# Quantitative intracellular retention of delivered RNAs through optimized cell fixation and immuno-staining

**DOI:** 10.1101/2021.07.06.451306

**Authors:** Prasath Paramasivam, Martin Stöter, Eloina Corradi, Irene Dalla Costa, Andreas Höijer, Stefano Bartesaghi, Alan Sabirsh, Lennart Lindfors, Marianna Yanez Arteta, Peter Nordberg, Shalini Andersson, Marie-Laure Baudet, Marc Bickle, Marino Zerial

## Abstract

Detection of nucleic acids within sub-cellular compartments is key to understanding their function. Determining the intracellular distribution of nucleic acids requires quantitative retention and estimation of their association with different organelles by immunofluorescence microscopy. This is important also for the delivery of nucleic acid therapeutics which depends on endocytic uptake and endosomal escape. However, the current methods fail to preserve the majority of exogenously delivered nucleic acids in the cytoplasm. To solve this problem, by monitoring Cy5-labeled mRNA delivered to primary human adipocytes via lipid nanoparticles (LNP), we optimized cell fixation, permeabilization and immuno-staining of a number of organelle markers, achieving quantitative retention of mRNA and allowing visualization of levels which escape detection using conventional procedures. Additionally, we demonstrated the protocol to be effective on exogenously delivered siRNA, miRNA, as well as endogenous miRNA. Our protocol is compatible with RNA probes of single molecule fluorescence in-situ hybridization (smFISH) and molecular beacon, thus demonstrating that it is broadly applicable to study a variety of nucleic acids.

## Introduction

The success of nucleic acid (e.g. siRNA, mRNA, anti-sense oligonucleotides) therapeutics depends on endocytosis and subsequent escape from endosomes to reach their site of action in the cytoplasm or the nucleus (Gilleron, Querbes et al. 2013, Kowalski, Rudra et al. 2019). Although endocytic uptake can be relatively specific and efficient, only a tiny fraction of endocytosed nucleic acids escapes from the endosomal lumen avoiding the ultimate degradation in lysosomes. The mechanism of nucleic acid therapeutics escape from endosomes is unclear (Dowdy 2017, Setten, Rossi et al. 2019). Therefore, to better understand the relationship between trafficking and escape as prerequisite to guide the development of delivery platforms, it is important to determine the subcellular distribution of nucleic acids in different endosomal compartments. This is not a trivial task because nucleic acids can enter cells via a range of uptake mechanisms (Gilleron, Querbes et al. 2013) and be transported to an endosomal network comprising several distinct endocytic compartments characterized by different transport kinetics, morphology and biochemical composition (Schmid, Sorkin et al. 2014, Dowdy 2017). A prerequisite for this analysis is to obtain quantitative estimates of the colocalization of nuclei acids to endosomal compartments labelled with specific markers by high resolution microscopy and image analysis (Gilleron, Querbes et al. 2013, Sahay, Querbes et al. 2013). The most common approach is to use fluorescently labeled nucleic acids or detect them by single molecule fluorescence in-situ hybridization (smFISH). The fluorescent nucleic acids can then be localized to endocytic compartments that are labelled by immunofluorescence staining (IFS) with specific antibodies. An alternative method is to image living cells expressing fluorescently tagged endosomal markers. However, this approach has two major drawbacks. First, cell lines expressing a panel of fluorescent endosomal markers covering the endocytic pathway are not always available. Second, it is not readily applicable to primary cells and tissues, especially human specimen. Consequently, IFS remains the simplest method for the intracellular characterization of nucleic acids. Nevertheless, the widely used method based on Formaldehyde (FA) for fixation and Triton X-100 for permeabilization fails to preserve nucleic acids quantitatively, especially for siRNA, miRNAs and mRNA (Urieli-Shoval, Meek et al. 1992, Pena, Sohn-Lee et al. 2009, Klopfleisch, Weiss et al. 2011, Fernández and Fuentes 2013).

Exogenous delivery of mRNA to cells has great potential for basic research but is also a major focus of nucleic acids-based therapeutics such as mRNA-based vaccines (Yanez Arteta, Kjellman et al. 2018, Kowalski, Rudra et al. 2019). Yet, only a handful of studies have addressed the sub-cellular trafficking of the delivered mRNA (Lorenz, Fotin-Mleczek et al. 2011, Kirschman, Bhosle et al. 2017). On the one hand, information on the efficiency of mRNA fixation and retention within the cytoplasm of the target cells using the standard IFS protocols is lacking. On the other hand, alternative strong fixatives, such as alcohols and glutaraldehyde, are often incompatible with antibody staining (Hopwood 1969, Farr and Nakane 1981, Hoetelmans, Prins et al. 2001). This led us to improve the methodology to retain the mRNA as well as other nucleic acids quantitatively and enable the characterization of its sub-cellular localization.

## Methods and materials

### Cell culture

HeLa cells were cultured in DMEM media complemented with 10% FBS Superior (Merck, S0615) and 50μg/mL Gentamycin (Gibco, G1397) at 37°C with 5% CO_2_. The day before a transfection 3000 HeLa cells in 50μL/well were seeded in 384 well plates using the drop dispenser (Multidrop, Thermo Fisher Scientific).

Human adipose stem cells (hASCs) from human subcutaneous White Adipose Tissue (WAT) was provided by AstraZeneca. hASCs were collected from patients undergoing elective surgery at Sahlgrenska University Hospital in Gothenburg, Sweden. All study subjects received written and oral information before giving written informed consent for the use of the tissue. The studies were approved by The Regional Ethical Review Board in Gothenburg, Sweden. All procedures performed in studies involving human participants were in accordance with the ethical standards of the institutional and national research committee and with the 1964 Helsinki declaration and its later amendments or comparable ethical standards. All subjects complied with ethical regulations. We adapted protocol from AstraZeneca (AZ) to differentiate hASCs to mature white-like adipocytes in 384 well format. Briefly, cryopreserved human adipose stem cells were resuspended in EGM-2 medium and centrifuged at 200xg for 5min. EGM-2 medium was prepared according to the manufacture’s protocol with EBM-2 medium supplemented with 5% FBS, all provided supplements, except hydrocortisone and GA-1000 (Lonza, Cat No. 3202, EGM^™^-2 MV BulletKit^™^ (CC-3156 & CC-41472). Cells were counted with a CASY cell counter (Schärfe System) and 4000 cells per well were seeded in 50μL EGM-2 medium containing 50 U/mL penicillin and 50 μg/mL streptomycin (P/S) (Gibco, 15140-122) into 384 well plates (Greiner Bio-One, 781092) using the drop dispenser Multidrop. The cells were incubated at 37°C and 5% CO_2_ for 3 to 4 days. For adipocyte differentiation, 90% confluent cells were incubated for 1 week with Basal Medium medium (Zenbio, BM-1) supplemented with 3% FBS Superior, 1μM dexamethasone (Sigma Aldrich), 500 μM 3-isobutyl-1-methyxanthine (Sigma Aldrich), 1μM pioglitazone (provided by AZ), P/S and 100 nM insulin (Actrapid Novonordisk, provided by AZ). Medium was replaced with BM-1 medium supplemented with 3% FBS Superior, 1μM dexamethasone, P/S and 100 nM insulin and cells were incubated for another 5 days. hASCs were tested and found free of mycoplasma.

### Chemicals

Formaldehyde (Merck), Triton X-100 (SERVA Electrophoresis GmbH), Digitonin (Sigma Aldrich) and Saponin (Sigma Aldrich), l-ethyl-3-(3-dimethylaminopropyl) carbodiimide (EDC) (Pierce^™^, Cat No. 22980), Disuccinimidyl suberate (DSS) (Pierce^™^, Cat No. 21555). FBS Superior (Merck, S0615), dexamethasone (Sigma Aldrich, D2915), 3-isobutyl-1-methyxanthine (Sigma Aldrich, I5879). Insulin (Actrapid Novonordisk) and Pioglitazone (AZ10080838) are provided by Astra Zeneca. Penicillin and Streptomycin (Gibco, 15140-122). LNPs were prepared with ((2-(dimethylamino)ethyl)azanediyl)bis(hexane-6,1-diyl)bis(2-hexyldecanoate) (AstraZeneca), cholesterol (Sigma Aldrich), 1,2-distearoyl-sn-glycero-3-phosphocholine (DSPC, CordenPharma), 1,2-dimyristoyl-sn-glycero-3-phosphoethanolamine-N [methoxy (polyethyleneglycol)-2000] (DMPE-PEG2000, NOF Corporation) and contained CleanCap® Enhanced Green Fluorescent Protein (eGFP) mRNA (5-methoxyuridine) (TriLink Biotechnologies, L-7201) and/or Clean Cap®Cyanine5 (Cy5) Enhanced Green Fluorescent Protein mRNA (5-methoxyuridine) (TriLink Biotechnologies, L-7701).

### LNP preparation and characterization

LNP were formulated by a bottom-up approach (Zhigaltsev, Belliveau et al. 2012) using a NanoAssemblr microfluidic apparatus (Precision NanoSystems Inc.). Prior to mixing, the lipids were dissolved in ethanol and mixed in the appropriate ratios while mRNA was diluted in RNase free 50 mM citrate buffer pH 3.0 (Teknova). The aqueous and ethanol solutions were mixed in a 3:1 volume ratio at a mixing rate of 12 mL/min to obtain LNP with a mRNA:lipid weight ratio of 10:1. Finally, they were dialyzed overnight using Slide-A-Lyzer G2 dialysis cassettes with a molecular weight cutoff of 10 K (Thermo Fisher Scientific). The size was determined by DLS measurements using a Zetasizer Nano ZS (Malvern Instruments Ltd.) The encapsulation and concentration of mRNA were determined using the Ribo-Green assay.

### LNP mRNA transfection

HeLa cells were transfected in the presence of 10% FBS Superior. Transfection for mature human white-like adipocytes was performed in the presence of fresh BM-1 medium supplemented with 1% human serum (Sigma, H4522). Both HeLa cells and mature human white like adipocytes were transfected with LNPs at a final mRNA concentration of 1.25μg/mL. LNP incubation times varied from 30min to 24h and is given for each experiment in figure legends.

### siRNA transfection

Alexa 647-labelled siRNAs were complexed with Interferrin (PolyPlus) for 10min in OptiMEM (siRNA final concentration = 10nM, Interferrin final volume/well/50µL = 0.1µL), added to HeLa cells in 10% FBS and incubated for 30min before washing and fixation. Fluorophore labeled siRNAs were ordered Sigma Aldrich from and sequence used are following. Sense strand: Alexa Fluor 647 – 5’-ACAUGAAGCAGCACGACUUdTdT-3’, Antisense strand: 5’-AAGUCGUGCUGCUUCAUGUdTdT-3’.

### Cell fixation

After LNP incubation, the cells were washed with PBS using the plate washer Power Washer 384 (PW384, Tecan). The cells were then fixed with formaldehyde using the drop dispenser (WellMate, Thermo Fisher Scientific) and washed 3 times with PBS. The percentage of formaldehyde (FA) and incubation time for experiments are given in figure legends. After FA fixation, an additional fixation with EDC for 2h was performed when required following previously reported procedure (Pena, Sohn-Lee et al. 2009). Briefly, the FA cells were washed with PBS and incubated with 0.2% glycine to quench remaining FA. The cells were incubated with methyl imidazole buffer 2 times each for 10min. The solution was replaced with freshly prepared EDC (0.16M EDC in 0.13M methyl imidazole buffer) and cells were incubated for 2h at room temperature. After EDC fixation the cells were incubated with 0.2% glycine for 5min and washed 3 times with PBS. DSS fixation was performed for 2h at room temperature following slightly modified manufacturer’s protocol (Thermo Fisher Scientific MAN0011240). The FA fixed cells were washed with PBS and replaced with 0.5mM DSS dissolved in DMSO. After 2h, the cells were washed 3 times with PBS. In some experiments, co-fixation of FA and DSS was used instead of sequential fixation for fast processing. Briefly, stock solution was prepared by adding FA directly from 37% to DSS DMSO solution. This stock was then directly added to cells to reach final concentration. Fixed cells were then washed, nuclei stained for imaging.

### Immuno-fluorescence staining

Cells were permeabilized with Triton X-100 or Saponin or Digitonin. The permeabilization conditions are described in figure legends. All detergents were added to cells using the liquid handling robot Biomek FX (Beckman Coulter) equipped with a 384 disposable tip head. After permeabilization, the cells were incubated with 3% BSA PBS blocking solution for 30min. Primary antibodies were incubated (in 3% BSA PBS solution) for either 1h (HeLa Kyoto cells) or 2h (Human primary white like adipocytes) to label endosomes. The cells were then washed 3 times and incubated with secondary antibodies (in 3% BSA PBS solution) for 1h. All blocking and antibody solutions were added using the liquid handling robot Fluent and washing steps were performed with Power Washer 384 (PW384, Tecan). After immunostaining the cells were incubated with DAPI (1µg/mL) and CMB (0.25µg/mL) to stain the cell nuclei and the cytoplasm respectively. All antibodies used in adipocytes and HeLa cells are listed in Supplementary Table 1.

### Single-molecule fluorescent in situ hybridization (smFISH) and immuno-fluorescence staining

smFISH was performed using reagents, including eGFP - CAL Fluor® Red 590 Dye probe and Transferrin receptor endogenous mRNA Alexa Fluor 647 probe, from Stellaris®. The protocol was compiled and modified from different manuals available from the manufacturer’s website (https://www.biosearchtech.com/support/resources/stellaris-protocols). Buffers were prepared according to the manufacturer, and volumes adapted to the 384 well plate format. In brief, after LNP incubation, the cells were fixed for 2h with 7.4% formaldehyde at RT. After washing with PBS, supernatant was removed manually with an 8-needle aspirator and 70% Ethanol was added for 1 h at 4°C. Under Digitonin permeabilization conditions, 0.004% Digitonin was added for 2min (adipocytes) or 0.001% 1min (HeLa cells). Note: Digitonin was first purified according to manufacturers’ protocol. (https://www.sigmaaldrich.com/content/dam/sigma-aldrich/docs/Sigma/Product_Information_Sheet/d5628pis.pdf). Since Digitonin is a natural extract subject to batch variation, we recommend either purifying a large batch sufficient for long-term use or optimizing the concentration for every single batch. After washing the cells with PBS using the plate washer PW384, cells were incubated in Wash Buffer A (40μL/well) for 2-5min. After removal of the supernatant the probe was added to the cells diluted 1:100 in Hybridization Buffer (12.5 μL/well, smFISH probe final concentration of 100nM) for about 16h at 37°C. The supernatant was removed and replaced twice with 40 μL Wash Buffer A for an incubation times of 30min at 37°C Finally, Wash Buffer A was replaced with Wash Buffer B for 2-5min. Finally Wash Buffer A was removed and DAPI (1µg/mL) and CMB (0.25µg/mL) in RNase-free PBS were used to stain the cell nuclei and the cytoplasm as described below. For sequential IFS and smFISH protocol, the fixed cells were first permeabilized with Digitonin, stained following the IFS protocol and fixed a second time with 3.7% FA for 10min at RT. Then the smFISH staining protocol was proceeded as described above with a second permeabilization with Ethanol. All antibodies used in this study are given in Supplementary Table 1.

### Fluorescence imaging and quantification

Where indicated, fixed cells were stained with 4′,6-diamidino-2-phenylindole (DAPI, 1μg/mL) and / or Cell Mask Blue (CMB, 0.5μg/mL). All imaging was performed on an automated spinning disc confocal microscope (Yokogawa CV7000) using a 60x 1.2NA objective. Live cell imaging was done at 37°C, 5% CO_2_ and humidified atmosphere. DAPI and CMB were acquired with a laser excitation at 405nm and an emission band pass filter BP445/45, GFP and Alexa 488 with a 488nm laser and BP525/50 filter, CAL Fluor® Red 590 with a 561nm laser and BP600/37 filter, Cy5 and Alexa 647 with a 640nm laser and a BP676/29 filter, and bright field using a halogen lamp. In most cases, 6 images were acquired per well and each condition was done in triplicate wells.

Image analysis was performed in custom designed software, MotionTracking (Collinet, Stöter et al. 2010). Images were first corrected for illumination, chromatic aberration and physical shift using multicolor beads. All fluorescent objects in corrected images were then segmented, their number and intensity per image mask area were calculated.

Image analysis was also done using Fiji (Schindelin, Arganda-Carreras et al. 2012) and CellProfiler (Carpenter, Jones et al. 2006) software. In brief, corrected images were pre-processed for segmentation in Fiji, LNP spots and nuclei were segmented and quantified in CellProfiler. Spot measurements for shape and intensities, as well as for spatial intensity distributions were loaded into KNIME data analysis software (Berthold, Cebron et al. 2008). Detected objects from auto-fluorescence were removed using a Random Forest classification algorithm. Data was visualized in KNIME and with customized R scripts.

### Xenopus laevis

*Xenopus laevis* embryos were obtained from *in vitro* fertilization, raised at 14-22°C in 0.1X MMR pH 7.5 and staged according to the table of Nieuwkoop and Faber (P.D. Nieuwkoop 1994). All animal experiments were approved by the Italian Minister of Health with the authorization no. 1159/2016-PR and no. 546/2017-PR according to art.31 of D.lgs. 26/2014.

### *Xenopus laevis* eye electroporation

Stage 26 embryos were anesthetized in 0.3 mg/mL MS222 (Sigma) in 1x MBS. The retinal primordium was injected with 0.5 µg/µl pCS2-CD63-GFP plasmid and 5 µM Molecular Beacon (MB) or 250 ng/μL pre-miRNA-181a-1, followed by electric pulses of 50ms duration at 1000ms intervals, delivered at 18V.

### *Xenopus laevis* organoculture

Cultures were performed as previously described (Bellon, Iyer et al. 2017). Briefly, glass-bottom dishes (MatTek) were first coated with poly-l-lysine (Sigma, 10 μg/mL 3 hours at 20°C) and then with Laminin (Sigma, 10 μg/mL 1 hour at 20°C). Eyes were dissected from stage 27 anesthetized embryos and cultured at 20°C for 40 h in 60% L-15 and 1% antibiotic-antimycotic (Thermo Fisher Scientific).

### Fixation and IFS in *Xenopus laevis* organoculture

Conventional protocol fixation and IFS was performed as follows. Organocultures were fixed in 2% PFA (Thermo Fisher Scientific), 7.5% sucrose (ACS reagent) for 30min and the fixed samples were permeabilized with 0.1% Triton X-100 (Fisher Chemical, in PBS 1x) 10min.

For improved fixation and IFS, Organocultures were fixed in 3.7% PFA (Thermo Fisher Scientific), 7.5% sucrose (ACS reagent) for 2 hours, washed thrice with PBS 1x (5min per wash), incubated in EtOH 70% (Sigma) for 1 hour at room temperature and washed 3 times again in PBS 1x. Cultures were permeabilized in 0.004% ethanol-purified Digitonin (Sigma) for 2min and blocked in filtered RNase free 3% BSA for 30min. Rabbit polyclonal anti-GFP (A-11122, Thermo Fisher Scientific, 1:500) or mouse anti-cy3 (sc-166894, Santa Cruz Biotechnology, 1:500) were used. Primary antibodies were diluted in 3% BSA and incubated at room temperature for two hours. Secondary antibodies Alexa Fluor AF-647 anti-rabbit (A-21246 Thermo Fisher Scientific, 1:1000) or anti-mouse (A-21237 Thermo Fisher Scientific, 1:1000) were incubated for 45min at room temperature. Cultures were washed thrice with PBS 1x and images were acquired without mounting as described below.

### Retinal ganglion cells (RGC) axons acquisition

Nikon Eclipse Ti2 inverted microscope equipped with Lumecor Spectra X LED light source and sCMOS camera (AndorZyla 4.2 Megapixel) was used with a Plan Apochromatic 60x/1.4 Oil objective. The acquisition mode was set to 12-bit and Gain4 (GFP and cy3 channel) and Gain 1 (cy5 channel) with a “Readout Rate” of 540 MHz and no binning. Exposure time and light intensity were kept invariant for the same batch of analysis. Representative images were adjusted for brightness and contrast.

### pre-miRNA *in vitro* synthesis and labeling

Pre-miR-181a-1 DNA template was obtained by two oligos (sequence below) after annealing and elongation at 12°C using 25U T4 DNA polymerase (NEB). The DNA template was purified with NucleoSpin PCR Clean-up kit (Macherey-Nagel). Pre-miR-181a-1 *in vitro* synthesis was performed using T7 MEGAshortscriptTM kit (Ambion) from 1 µg purified DNA and following manufacturer’s instructions, including template removal by DNase TURBO digestion. 5 µg pre-miR-181a-1 were labeled with cy3 using Nucleic Acid Labeling Kit (Mirus) following the manufacturer’s instructions.

Pre-miR-181a-1_T7_Fw: 5’-TAATACGACTCACTATAGAACATTCAACGCTGTCGGTGAGTTTGGTATCTAAAGGC-3’

Pre-miR-181a-1_Rv: 5’-TGTACAGTCAACGATCGATGGTTTGCCTTTAGATACCAAACTCACCG-3’

### Oligonucleotides and plasmid

cy3-BHQ2 pre-miR-181a-1 molecular beacon (Eurogentec): 5’-CAUUGC**C**UUUA**G**AUACCAAUG -3’, 2’-O-methyl ribose backbone and 2 LNA nucleotides (bold)(Corradi, Dalla Costa et al. 2020).Plasmid, pCS2-CD63-eGFP (Corradi, Dalla Costa et al. 2020).

## Results and discussion

### Significant loss of exogenously delivered mRNA after fixation and permeabilization

Several studies reported loss of endogenous mRNA from cells during commonly used IFS protocols (Urieli-Shoval, Meek et al. 1992, Pena, Sohn-Lee et al. 2009, Klopfleisch, Weiss et al. 2011, Sylwestrak, Rajasethupathy et al. 2016). Exogenously delivered nucleic acids internalized in endosomal organelles must be preserved by fixation and permeabilization methods. To address this problem, we used mRNA labelled with the Cy5 fluorophore (Cy5 mRNA), formulated in lipid nanoparticles (LNP) and delivered to human primary adipocytes, as clinically relevant cell system (Yanez Arteta, Kjellman et al. 2018). LNP subcutaneous administration is aimed to reach the fatty layer beneath the epidermis and dermis of the skin. Primary human adipocytes are therefore the most relevant cell model to study uptake of drugs administered by this route. First, we estimated the extent of signal loss by quantifying the efficiency of mRNA fixation using a widely used formaldehyde-based protocol (3.7% for 10min at RT). Although LNP Cy5 mRNA were detectable in living cells as soon as 30min of incubation (Figure 1A), only very low signal was retained in fixed cells, suggesting that the fixation protocol fails to quantitatively retain mRNA in the cytoplasm. Second, we quantified the loss of mRNA upon fixation and permeabilization with detergent. For this, we incubated LNP Cy5 mRNA with adipocytes for 24h to accumulate high levels of mRNA and maximize the signal. The cells were then fixed and permeabilized with Triton X-100. Strikingly, we estimated a loss of 83.5% ± 0.5 of Cy5 mRNA signal in cells subjected to permeabilization compared with control cells, i.e. fixed and without permeabilization (Figure 1B & C). Such a loss was not limited to mRNA, as it was also observed with siRNAs delivered via commercial transfection reagents in HeLa cells permeabilized with Triton X-100 (Figure 1D & E). Overall, these results suggest that loss of exogenously delivered nucleic acid occurs in both fixation and permeabilization steps. These results prompted us to improve the protocol for better retention of nucleic acids in cells.

**Figure 1:**
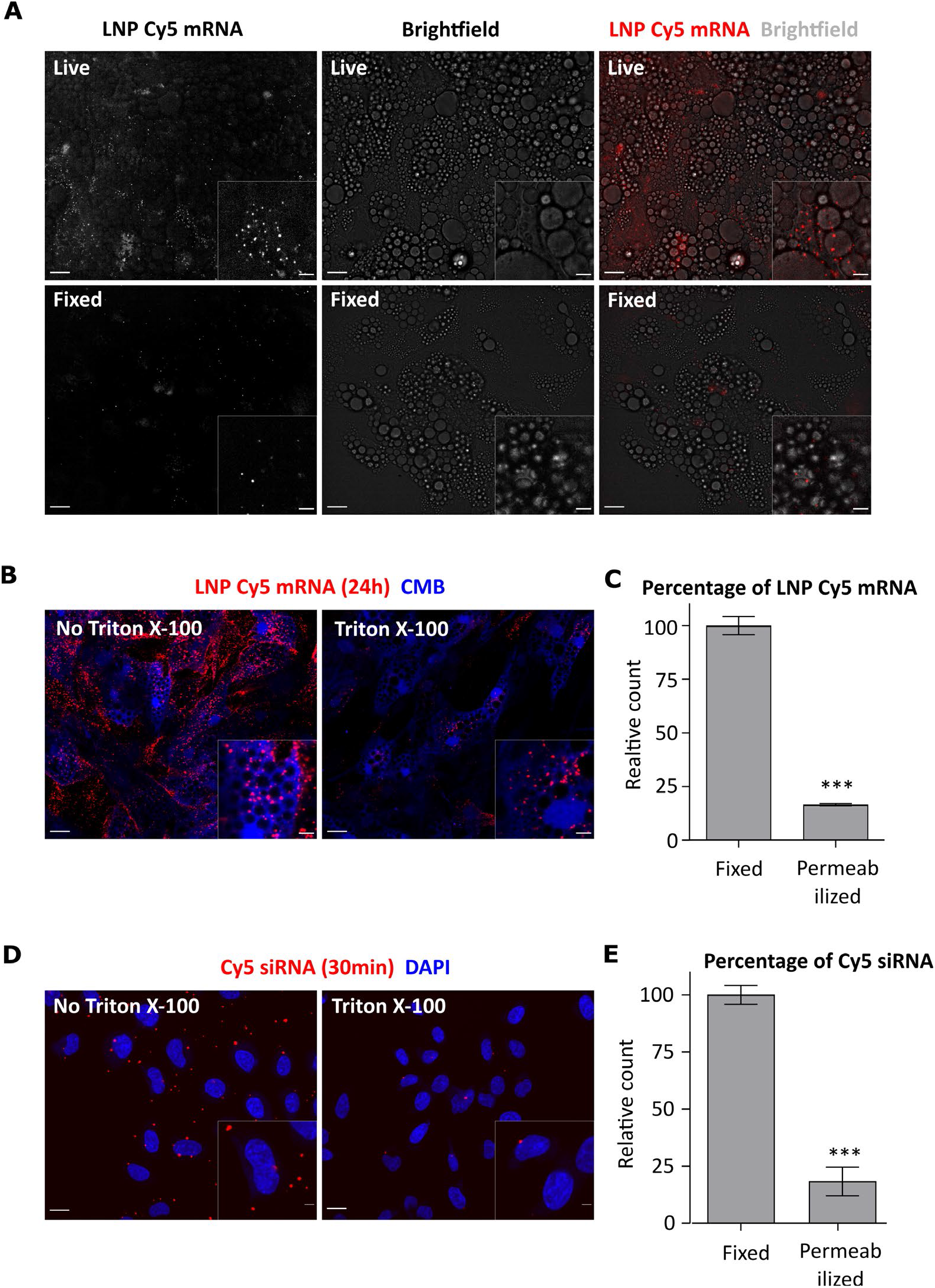
Poor Cy5 mRNA retention during cell fixation and permeabilization. (**A**) LNP Cy5 mRNA uptake (30min) in human primary adipocytes. The amount of Cy5 mRNA signal in FA-fixed cells (3.7% for 10min) is very low compared to living cells. The circular structures in the bright field images are lipid droplets of adipocytes. (**B**) Representative images of cells incubated with LNP Cy5 mRNA (24h) that were either only fixed or fixed and permeabilized with Triton X-100 (0.1%, 10min). The dark holes in the cytoplasm correspond to lipid droplets that cannot be stained with CMB. (**C**) The quantification shows that the percentage of Cy5 mRNA object count is significantly lower in permeabilized cells compared to non-permeabilized cells. (*** = p<0.0001 relative to “Fixed”). All conditions were performed in triplicates, Mean ± SEM). These data were taken from the experiments of Figure 3. (**D**) HeLa cells incubated with the commercial transfection reagent Interferrin were either fixed or fixed and permeabilized. Representative images show loss of Cy5-siRNA in fixed cells after Triton X-100 (0.1%, 10min) permeabilization. (**E**) Quantification of Cy5 siRNA show 81.7% ± 6.2 loss of siRNAs in Triton-X 100 permeabilized cells compared only fixed cells. These data were taken from experiments of Figure 5B. N = 3 independent experiments. Mean ± SEM are displayed. The scale bars of full images are 20µm, inset images 5µm.

### Formaldehyde concentration and incubation time are crucial to retain mRNA quantitatively

Previous studies have shown that after FA fixation, an additional fixation with 1-Ethyl-3-(3-Dimethylaminopropyl) Carbodiimide (EDC) or Disuccinimidyl suberate (DSS) can covalently fix nucleic acids in tissues (Pena, Sohn-Lee et al. 2009, Sylwestrak, Rajasethupathy et al. 2016). We tested these fixatives first in HeLa cells as they are easier to culture and process than primary adipocytes. To determine whether fixation with EDC and DSS is compatible with IFS of endosomal compartments, we used anti EEA1 antibodies to label early endosomes. Although EDC and DSS fixation following FA improved the retention of Cy5 mRNA, a significant loss of signal persisted (Figure 2A & B). Moreover, EDC treatment was incompatible with IFS as it increased the background fluorescence upon staining with EEA1 antibodies (Figure 2C), making colocalization-based analysis impossible. In contrast, DSS significantly reduced the EEA1 signal in IFS. These results indicate that fixation with EDC and DSS does improve the retention of mRNA signal but is not compatible with IFS.

**Figure 2:**
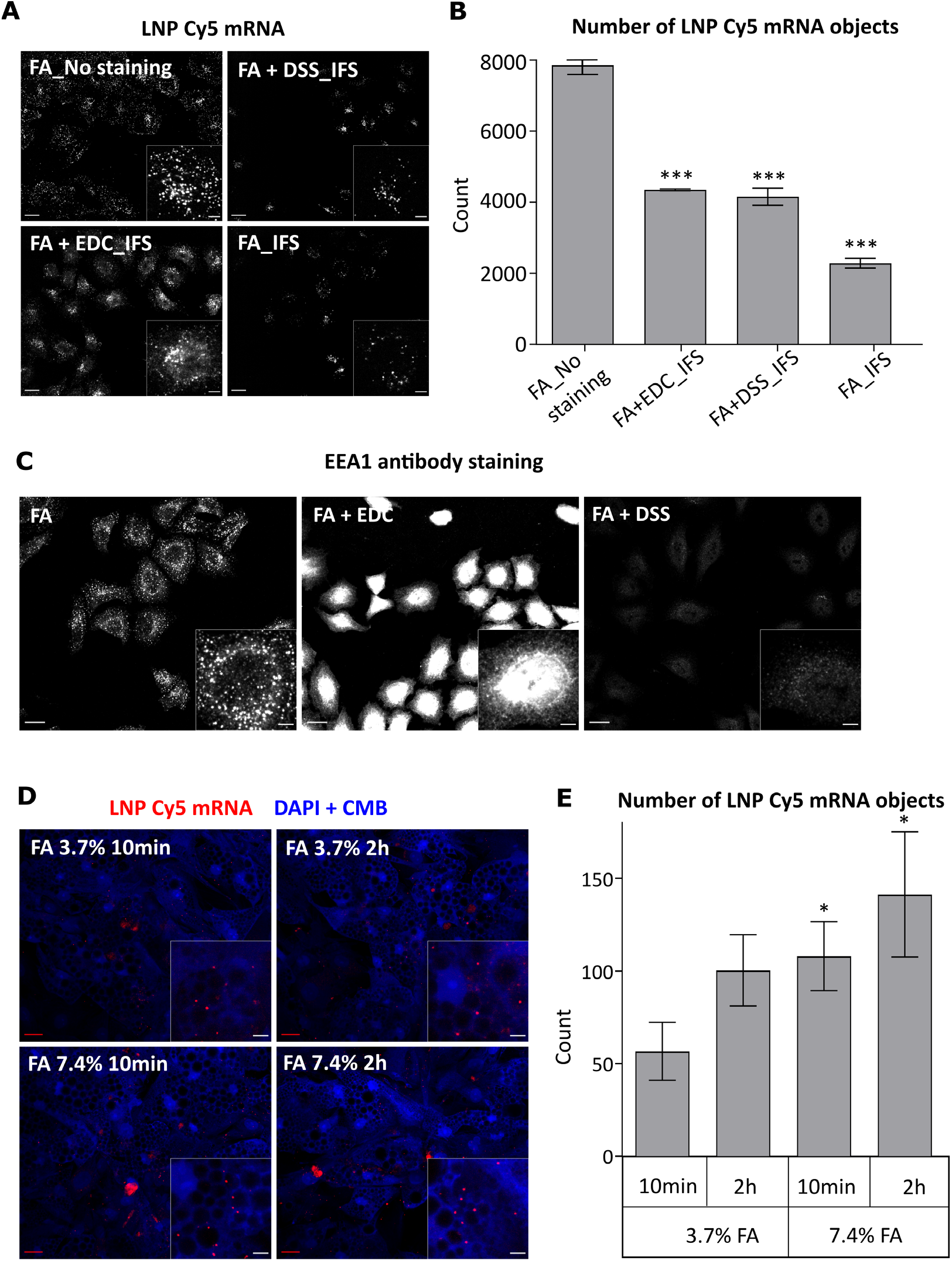
Higher FA concentration and incubation time retain more Cy5 mRNA in cells. (**A**) LNP Cy5 mRNA retention in EDC and DSS fixed HeLa cells. Cells incubated with LNP Cy5 mRNA (2h) were either fixed with 3.7% FA (10min) alone or fixed additionally with either EDC or DSS (2h). The images were taken after Triton X-100 permeabilization and immuno-staining with EEA1 antibodies. (**B**) Quantification of Cy5 mRNA retention shows that after permeabilization, both EDC and DSS fixation retain more mRNA compared to cell fixed with FA alone. All conditions were performed in duplicates. (*** p<0.0001 relative to “FA No staining”). (**C**) Representative images of EEA1 antibody staining. The images show artifacts upon EDC fixation and poor staining in DSS fixed conditions, indicating incompatibility with IFS. (**D**) LNP Cy5 mRNA retention after FA fixation in human primary adipocytes. Cells incubated with LNP Cy5 mRNA (30min) were fixed with FA at concentration and incubation time as indicated. (**E**) Quantification shows improved Cy5 mRNA retention with increasing FA concentration and incubation time. (* p<0.04 relative to “FA_3.7% FA_10min”). All conditions were performed in triplicates. Mean ± SEM. The scale bars of full images are 20µm and inset images are 5µm.

We therefore returned to formaldehyde-based fixation as it is a method widely applied to IFS. We tested formaldehyde at higher concentration and longer incubation times in order to improve the crosslinking of proteins and thus trap mRNA better in HeLa cells and human primary adipocytes. Our results show that higher FA concentration (7.4%) and longer incubation time (2h) retained more signal compared to control (3.7% FA 10min incubation) (Figure 2D & E). Similar results were obtained in HeLa cells (Supplementary Figure 1A). Additionally, FA with longer incubation time and higher concentration retained comparable amount of mRNA signal to FA and DSS co-fixed condition (Supplementary Figure 1A & B). Thus, the simple modified FA fixative method is sufficient to fix mRNA better and requirement of special mRNA fixatives are not necessary for exogenously delivered mRNA.

### Mild permeabilization method is crucial to retain more mRNA during IFS

A significant loss of Cy5 mRNA signal occurred during permeabilization of cells (Figure 1B). Presumably, Triton X-100 solubilizes the endosomal membrane and causes the subsequent loss of mRNA signal that are insufficiently fixed. Saponin and Digitonin are mild detergents that interact with cholesterol and form pores on the plasma membrane but do not efficiently permeabilize the endosomal membrane (Malerød, Stuffers et al. 2007, Sudji, Subburaj et al. 2015). Such permeabilization would be sufficient for antibodies to pass through the plasma membrane during IFS but prevent the loss of mRNA from the endosomes. We first tested previously reported conditions of Saponin and Digitonin detergents for permeabilization in HeLa cells (Villaseñor, Nonaka et al. 2015), in a range of concentrations below the critical micelle concentration (CMC) (Saponin CMC = 0.5g/L - 0.8g/L, Digitonin CMC = 0.25–0.5 mM). Compared to non-permeabilized cells, Digitonin retained a large proportion (93.56% ± 2.48) of Cy5 mRNA signal in contrast to Triton X-100 which retained only a minor fraction (9.18% ± 1.08) (Figure 3A & Supplementary Figure 2A). Digitonin permeabilization under this condition was also compatible with IFS as the EEA1 antibodies signal was comparable to classical Triton X-100 permeabilization conditions (Figure 3B & Supplementary Figure B & D). Interestingly, permeabilization with Saponin reduced the Cy5 mRNA signal to 12.5% ± 0.54. Both Triton X-100 and Saponin showed no improvement of Cy5 mRNA signal at lower concentrations (Supplementary Figure 2C).

**Figure 3:**
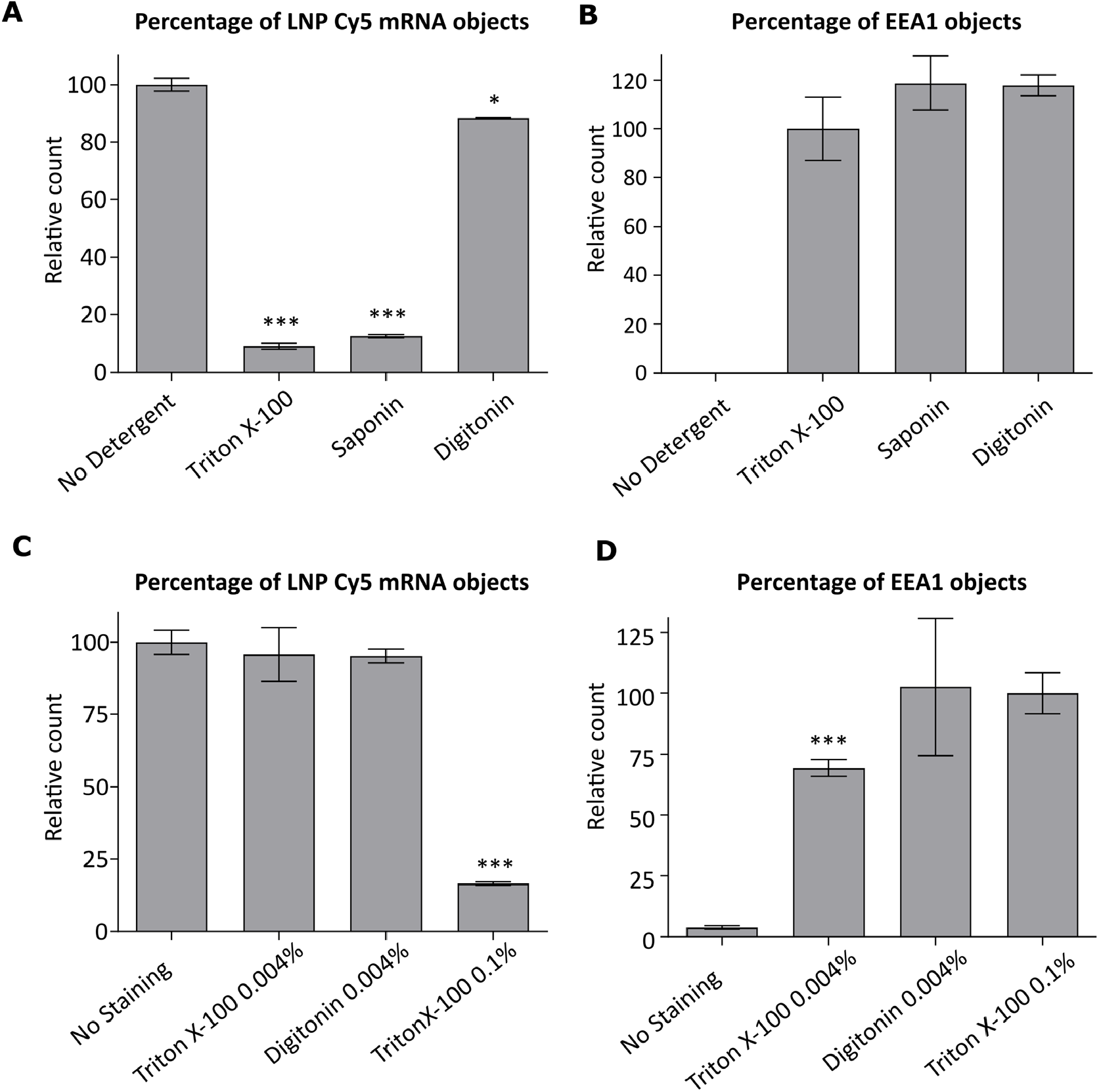
Mild cell permeabilization with Digitonin prevent loss of Cy5 mRNA. (**A**) Cy5 mRNA retention after permeabilization with detergents. HeLa cells treated with LNP Cy5 mRNA (1h) are FA fixed (FA 3.7%, 10min), permeabilized either with Triton X-100 (0.1%, 10min) or Saponin (0.1%, 10min) or Digitonin (0.001%, 1min), and immuno-stained with EEA1 antibodies. Compared to non permeabilized cells (no staining), both Triton X-100 and Saponin treatment show significant mRNA loss, whereas Digitonin retains mRNA signal. (* p =0.053, *** p <0.0001 relative to “No Detergent”) (**B**) The graph shows that EEA1 staining is good with all detergent permeabilization conditions (in quadruplicates). (**C**) Optimization of Digitonin permeabilization in human primary adipocytes. Cells treated with LNP Cy5 mRNA (24h) were fixed with FA (3.7% for 10min) and permeabilized with Digitonin or Triton X-100 at the indicated concentrations. All conditions were done in triplicates. The graph show that Digitonin and Triton X-100 retain most mRNAs at 0.004% concentration and 1min incubation compared to classical method (0.1% Triton X-100, 10min). (*** p <0.0001 relative to “No staining”). (**D**) Quantifications of EEA1 objects show that IFS in Digitonin permeabilized condition is comparable to the classical method. (*** p <0.0008 relative to “Triton X-100 0.1%”, Non-significant and No staining condition p values are not displayed). Mean ± SEM.

Since Digitonin permeabilization did not lead to significant loss of mRNA and was compatible with IFS in HeLa cells, we proceeded to test the protocol in human primary adipocytes. Given the abundance of lipid droplets in adipocytes, we first optimized the Digitonin permeabilization step by testing various concentrations and incubation times on Cy5 mRNA retention and EEA1 antibodies staining in comparison to Triton X-100 permeabilization. In adipocytes, 0.004% Digitonin permeabilization for 1min retained 95.21% ± 2.38 of mRNA signal and gave EEA1 staining comparable to that achieved with classical 0.1% Triton X-100 10min permeabilization (Figure 3C-D, Supplementary Figure 3A-B). Interestingly, the mRNA signal loss did not depend on the Digitonin incubation time (Supplementary Figure 4A). In contrast, permeabilization with Triton X-100 yielded sub-optimal results at all concentrations tested. Longer permeabilization times with Triton X-100 caused a significant loss of mRNA signal and reducing the concentration retained the mRNA signal but also reduced the EEA1 staining (Supplementary Figure 4A and B). When cells were permeabilized with 0.004% Triton X-100 for 1min, 95.72% ± 9.29 of mRNA signal was retained. However, the above mentioned condition showed less IFS efficiency compared to the classical protocol (0.1% Triton X-100 for 10min) (Figure 3D, Supplementary Figure 3B). Altogether, these results suggest that permeabilization with Digitonin retains most mRNA signal across various concentrations and is compatible with IFS in adipocytes.

We also found two other steps that caused major nucleic acids mRNA loss. First, in contrast to DAPI, nuclear staining with Hoechst quenched ∼ 80% of the mRNA signal (Supplementary Figure 5A). Second, BSA blocking caused mRNA loss presumably due to RNase contaminants (Data not shown). Although usage of these reagents is not necessarily common, we recommend to avoid Hoechst or use the lowest possible concentration for fluorescence microscopy based mRNA studies and take precautions such as including RNase inhibitor while using BSA blocking solutions.

### Adaptation of improved fixation and permeabilization methods for smFISH

The optimized protocol was established using LNP delivered Cy5-mRNA. To demonstrate that the protocol is applicable to unlabeled mRNA, we tested its compatibility with smFISH, as this method is widely applied to visualize endogenous mRNA. As a prerequisite, we first verified whether the smFISH probe can efficiently detect the majority of mRNAs under our optimized protocol, by quantifying the co-localization of smFISH probe and pre-labeled Cy5-mRNA. To this end, we first performed smFISH staining on LNP Cy5 mRNA deposited on glass slides. About 98.09% ± 0.48 of Cy5 mRNA were also labeled with the smFISH probe (eGFP - CAL Fluor® Red 590 Dye) (Supplementary Figure 6), demonstrating that it can be used to detect the mRNA. Next, to determine whether the smFISH staining can be applied under the improved fixation and permeabilization protocol, we incubated Cy5 mRNA in adipocytes for 30min, fixed and permeabilized the cells as described in the previous section. We also used ethanol (EtOH) permeabilization as recommended by the smFISH probe manufacturer for comparison under our fixation condition. As shown in Figure 4B, EtOH permeabilization labelled 99.19% ± 0.02 (in HeLa cells 97.59% ± 0.25, Supplementary Figure 8A) of Cy5 mRNA with the smFISH-570 probe, suggesting that 2h fixation with 7.4% FA did not affect the smFISH (Figure 4A & B). We also noted that the Cy5 mRNA signal was not lost after EtOH permeabilization compared to non-permeabilization control conditions (Figure 4C). Nonetheless, EtOH permeabilization showed insufficient labelling or produced artifacts with some antibodies tested (Supplementary Figure 7). These results point at incompatibility between EtOH permeabilization and IFS. In contrast, Digitonin permeabilization resulted in only 81.69% ± 0.22 (in HeLa cells 74.5% ± 4.36, Supplementary Figure 8B) of Cy5 mRNA labeled with the smFISH-570 probe. Overall, these results suggest that 1) higher FA concentration and longer fixation times do not affect smFISH efficiency, 2) both EtOH and Digitonin permeabilization methods retain most mRNA signal but only Digitonin is compatible with IFS, 3) despite retaining mRNA, Digitonin permeabilization alone is insufficient for smFISH mRNA labelling.

**Figure 4:**
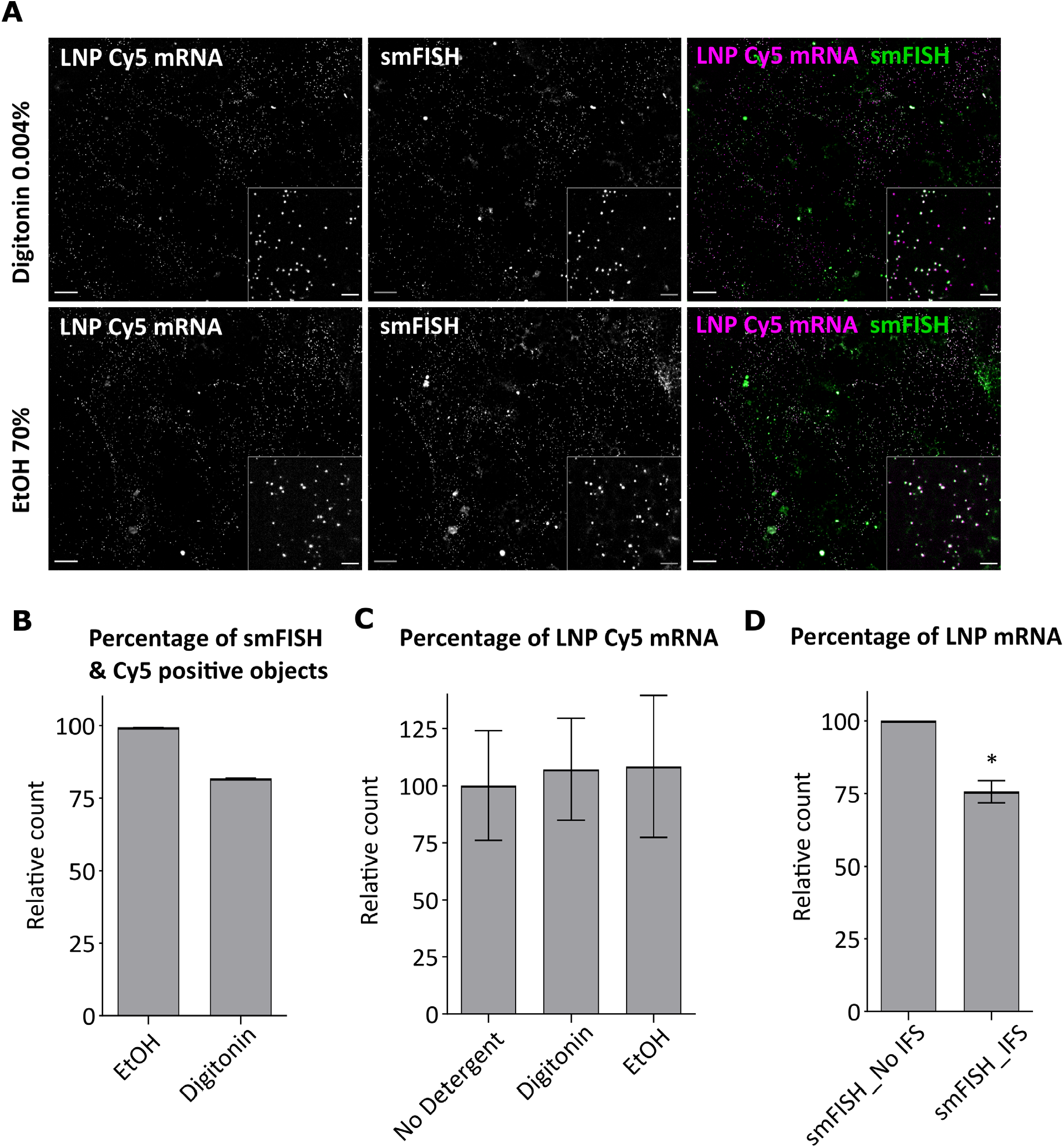
Improved fixation and permeabilization methods can be adapted for smFISH. (**A**) smFISH was performed in adipocytes incubated with LNP Cy5 mRNA after fixation (7.4% FA, 2h) and permeabilization with ethanol or Digitonin. The images show that a subset of Cy5 mRNA are not labeled for smFISH in Digitonin permeabilized cells. All conditions were done in triplicates (**B**) The graph show that the smFISH labelling of Cy5 mRNA is nearly 100% with ethanol, whereas a subset of Cy5 mRNA objects are not labelled in Digitonin permeabilized cells. (**C**) The graph shows retention of the Cy5 mRNA after EtOH and Digitonin permeabilization. (**D**) Adipocytes incubated with LNP unlabeled mRNA (1h) was stained as described in the result section. The graph illustrates that mRNA is retained after IFS. (* p=0.04 relative to smFISH_No IFS). N= 3 independent experiments, mean ± SEM.

To solve this problem, we designed a two-step permeabilization protocol for IFS and detected unlabeled mRNA using smFISH. First, after 7.4% 2h FA fixation, we performed Digitonin permeabilization and immuno-staining. Second, the cells were fixed again with classical fixation (3.7% FA for 10min) to preserve the antibody signal after IFS. Third, we performed EtOH permeabilization and smFISH staining to label the mRNA. As shown in Figure 4D, the protocol retained about 75.69% ± 3.84 of mRNA signal and was also compatible with IFS (Supplementary Figure 9). It is important to note that neither permeabilization with Digitonin nor with EtOH caused a significant loss of mRNA (Figure 4C) but this occurred only upon IFS and smFISH (25% loss in adipocytes, 38% in HeLa cells, Figure 4D & Supplementary Figure 8C respectively).

Next, we tested whether our improved smFISH protocol is generally compatible with IFS of cytoplasmic organelles. For this, we tested a set of 8 antibodies against various markers of organelles of the endocytic pathway, such as APPL1, Rab5, Rab11, ANKFY1, LBPA, LAMP1, CAV1 and LC3. We found that the protocol consistently retained up to 95% of mRNA without compromising the quality of organelle marker staining (Supplementary Figure 10I –J and Supplementary Figure 11A). Altogether, these results show that our modified IFS compatible protocol can be combined with smFISH and preserve a significantly higher amount of mRNA than the current protocol.

### Adaptation of improved protocol for other nucleic acids

Finally, we wanted to determine whether our protocol is generally applicable to different RNAs, both endogenous and exogenously delivered. We first tested it on the retention of endogenous mRNA in HeLa cells, focusing on the Transferrin receptor (TFR) mRNA, since it is expressed in a wide range of cells. We noted no TFR mRNA loss by smFISH after Triton-X 100 permeabilization compared to Digitonin permeabilized control (Supplementary Figure 12). Therefore, the problem of lack of retention applies to exogenously delivered mRNA or siRNA but not endogenous cytoplasmic mRNA. To determine whether our method could improve the retention of exogenously delivered siRNA in HeLa cells, we tested the efficacy of the fixation and permeabilization steps. Interestingly, increased FA concentration and longer incubation time did not improve retention of siRNA (Figure 5A). One possibility is that small nucleic acids like siRNAs (usually 22 nucleotide base pairs) have a lower probability to be cross-linked and trapped intracellularly upon fixation than large mRNA molecules (several hundred nucleotide bases long). However, longer fixation times helped to retain more siRNAs during the permeabilization step (Figure 5B, compare 64.9% ± 8 siRNA retention in 10min FA fixed cells Vs 84.7% ± 7 in 2h FA fixed cells under Digitonin permeabilization condition). In addition, we noted that 3.7% 2h FA fixation, sufficiently retained siRNA with good EEA1 staining but 7.4% fixation reduced EEA1 staining in HeLa cells (Supplementary Figure 13). Therefore, we recommend to optimize fixative concentration in each cell model system to ensure minimum loss of siRNAs while preserving the signal by antibody staining. Moreover, similar to mRNA (Supplementary Figure 5A), we recommend to avoid or use lowest possible concentration of Hoechst nuclear staining as it quenched ∼ 55% of siRNA signal (Supplementary Figure 6B).

**Figure 5:**
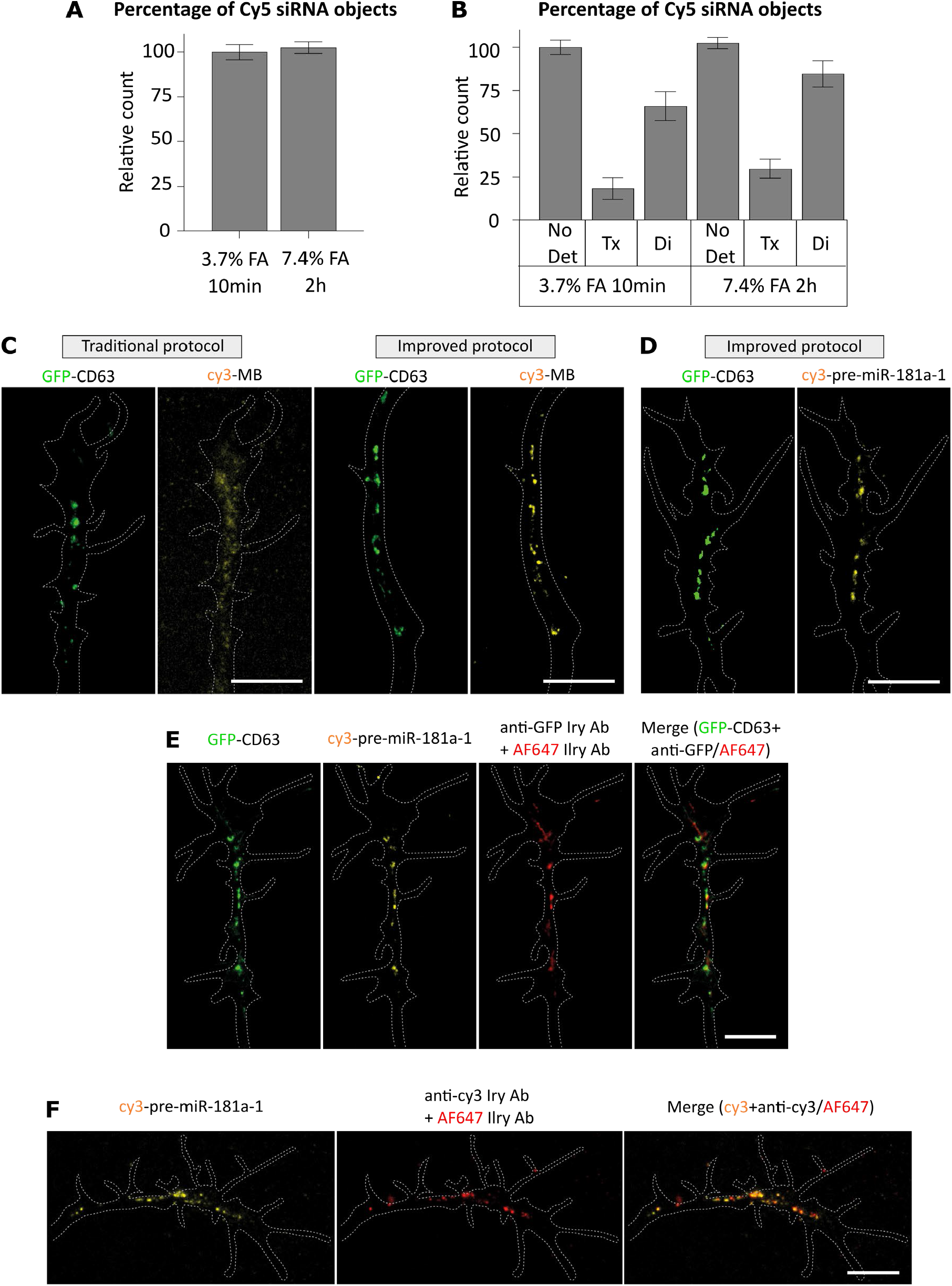
Improved intracellular retention of exogenously delivered siRNA and endogenous miRNA. (**A**) Cy5 siRNA + Interferrin incubated in HeLa cells for 30min were fixed with FA as indicated in the graphs. In contrast to mRNA fixation data shown in Supplementary Figure 1, fixation with 7.4% FA for 2h does not improve siRNA retention (**B**) The graph illustrates siRNA retention after 3.7 and 7.4% FA fixation (10min) under Digitonin (0.002%, 2min) and Triton-X (0.1%, 10min) permeabilization. N = 3 independent experiments. Mean ± SEM are displayed. (**C**) Comparison of miRNA retention using traditional and improved protocol in *Xenopus laevis* organoculture. Concentrations used are as follows: 5 μM MB; 250 ng/μL cy3-pre-miR-181a-1; 0.5 μg/μL pCS2-CD63-eGFP (see Materials & Methods). (**D, E & F**) Representative images. Dashed white lines delineate axons. Compare to traditional protocol, endogenous (pre)-miRNA retention improved significantly with our optimized protocol. 54 axons were analyzed in total from up to 7 independent experiments. Experimental details are described in the methods and main text. Abbreviations: CD63-eGFP, CD63-eGFP expressing pCS2 plasmid; PFA, paraformaldehyde. Scale bars: 10 μm.

Since the retention of siRNAs was improved by our protocol, we tested its effect on other forms of small nucleic acids. Endogenous microRNAs (miRNAs) can associate with late endosomes/lysosomes (LE/Ly) and hitchhike them for intracellular transport (Gibbings, Ciaudo et al. 2009, Lee, Pressman et al. 2009, Corradi and Baudet 2020, Corradi, Dalla Costa et al. 2020). Therefore, we visualized miRNAs docked to LE/Ly by two strategies. First, we detected endogenous precursor miRNAs (pre-miRNAs) via cy3-labeled molecular beacon (MB) (Supplementary Figure 14A) (Corradi, Dalla Costa et al. 2020) and assessed its colocalisation with LE/Ly marker transmembrane tetraspanin CD63-GFP (Pols and Klumperman 2009, Corradi, Dalla Costa et al. 2020) in elongating axons from *Xenopus laevis* whole eye explant culture (Supplementary Figure 14B). Whilst the standard protocol yielded a diffuse pre-miRNA-associated signal (Fig. 6C), with our optimized protocol the signal was significantly improved and appeared as discrete puncta colocalising with CD63-GFP marked vesicles (Figure 5C), consistent with previous findings in living cells (Corradi, Dalla Costa et al. 2020). Second, we investigated exogenously delivered (by electroporation) synthetic cy3-labeled pre-miRNA (Supplementary Figure 14A & B). Again, we detected discrete puncta (Figure 5D) as in live cells (Corradi, Dalla Costa et al. 2020), suggesting that our protocol dramatically improves the retention of both endogenous and exogenous pre-miRNAs. Finally, we tested whether our miRNA detection protocol is fully compatible with immunocytochemistry (Figure 5E). Anti-GFP and anti-cy3 readily detected CD63-GFP (Figure 5E) and cy3-labeled-pre-miRNA (Figure 5F), respectively, without altering miRNA retention.

In conclusion, in this study we highlighted the importance of verifying the loss of exogenously delivered and endogenous nucleic acids in cells subjected to IFS using the standard fixation and permeabilization protocol. The standard protocol allows the retention of only a minor fraction of signal. Lack of quantitative retention of mRNA can skew the subcellular mRNA distribution results yielding misleading conclusions. We outlined an improved fixation and IFS methodology that can retain quantitative amounts of exogenously delivered mRNA, siRNA and miRNA, as well as endogenous miRNA in the cell cytoplasm. This methodology is especially critical when the intracellular levels of RNA are very low. Our optimized protocol was applied to adipocytes, HeLa cells and axons from *Xenopus laevis* whole eye explant culture, suggesting it is generally applicable with minor modifications depending on the specific cell type. Our study can guide and offer appropriate solutions to researchers who study nucleic acids in various cell model systems to focus on critical steps, save time, resources as well as enable studies that were previously not feasible due to significant nucleic acid loss (similar to miRNA detection in endosome trafficking in axons). Since our methodology is compatible with probes of smFISH and molecular beacon, we hope it will be broadly applied to fluorescence-based quantification studies of various forms of exogenously delivered and endogenous nucleic acids.

## Acknowledgements

We thank Dr. Yannis Kalaidzidis for helpful discussions and suggestions. We would also like to thank the following Services and Facilities of the Max Planck Institute of Molecular Cell Biology and Genetics for their support: Light Microscopy Facility (LMF) and the High-Throughput Technology Development Studio (HT-TDS) Facility. We thank the Center for Information Services and High Performance Computing (ZIH) of the TU Dresden for the generous provision of computing power. This work was financially supported by the Max Planck Society (MPG) and Astra Zeneca. E.C, I.D.C and M-LB also thank Marie Curie Career Integration (618969 GUIDANCE-miR), G. Armenise-Harvard Foundation, MIUR SIR (RBSI144NZ4) and MIUR PRIN 2017 (2017A9MK4R) grants (to M.-L.B.) for funding this research.

## Author contributions

M.Z, P.P, M.S, and M.B, designed the experiments. M.Y.A formulated LNPs. P.P and M.S performed experiments in cells. M.S. performed automated image acquisitions. E.C, I.D.C and M-LB designed analysed and interpreted experiments in miRNAs. E.C, and M-LB wrote the miRNA part of the manuscript. A.H, S.B, A.S, L.L, P.N and S.A commented on results and interpretations. M.Z and P.P wrote the manuscript with edits by M.S, M.B, and M.Y.A.

