## Supplementary Figures for "Quantitative intracellular retention of delivered RNAs through optimized cell fixation and immuno-staining"

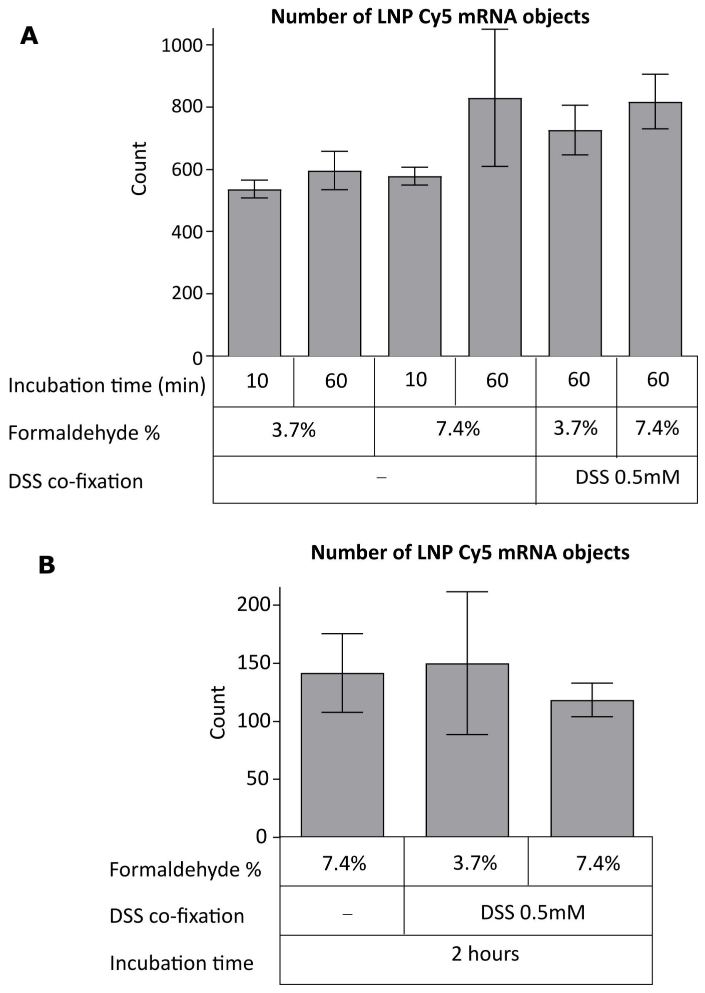


**Supplementary Figure 1: LNP Cy5 mRNA retention in FA and DSS fixed HeLa cells (A) and human primary adipocytes (B).** (**A**) Cells incubated with LNP Cy5 mRNA were either fixed with FA alone or FA and DSS co-fixation at a concentration and incubation time as indicated. Quantification shows that higher FA concentration and incubation time retains Cy5 mRNA comparable to FA and DSS co-fixation method. All conditions were done in quadruplicates. (**B**) Quantification of Cy5 mRNA retention in human primary adipocytes. The graph shows that 7.4% FA, 2h fixation without DSS is comparable to FA and DSS co-fixation method. All conditions were performed in triplicates. Mean ± SEM are displayed. FA 7.4%, 2h fixation condition in this graph is also presented in Figure 2 E.

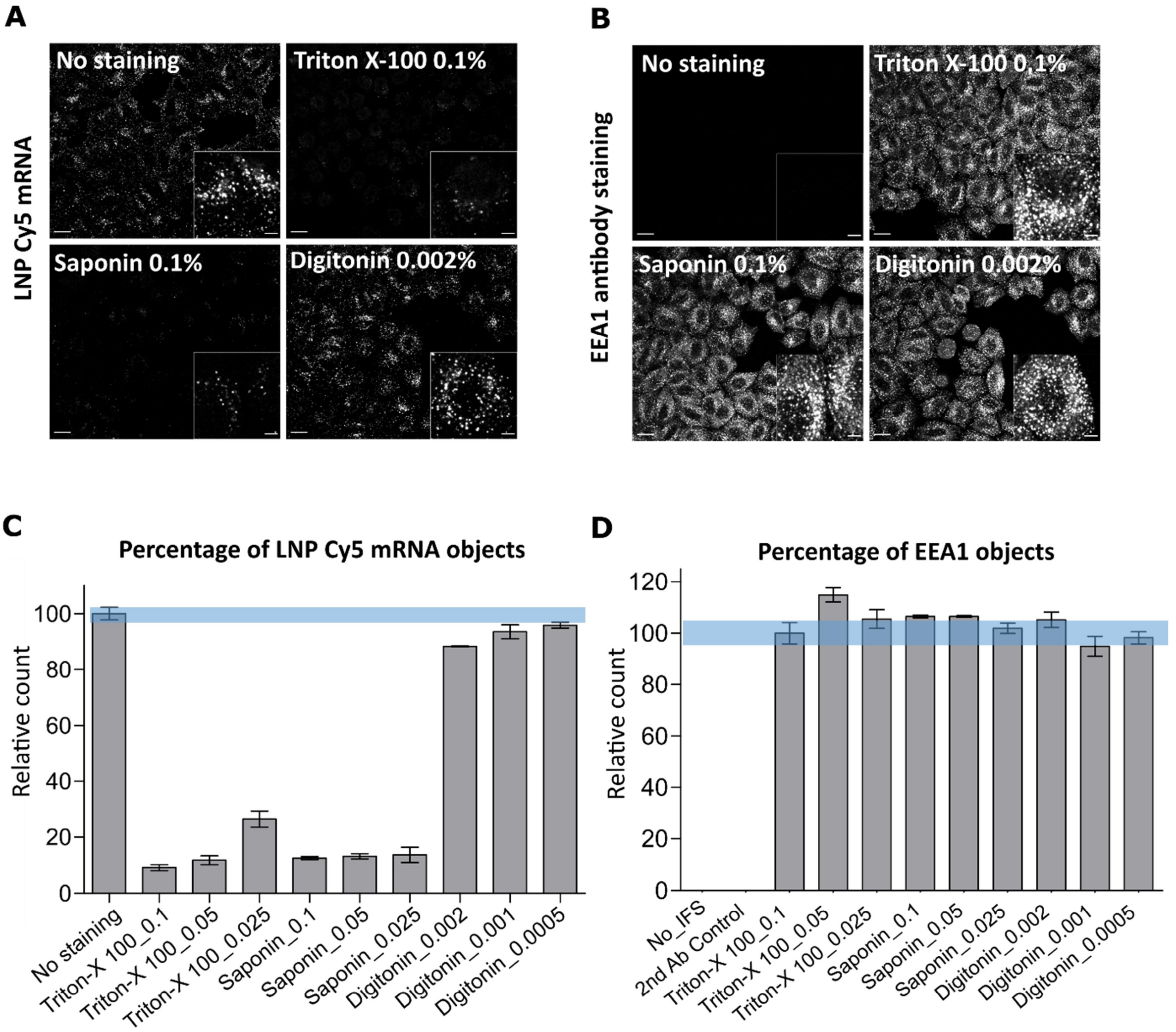


**Supplementary Figure 2: mRNA retention in HeLa cells under various detergent concentrations.** Experimental details and few selected conditions are described in Figure 3A. (**A**) Representative images of LNP Cy5 mRNA signal presented in Figure 3A. (**B**) Representative images of EEA1 antibody staining presented in Figure 3B. (**C**) LNP Cy5 mRNA quantification for all tested conditions. The graph shows that Digitonin permeabilization retains most mRNA signal. (**D**) Quantification of EEA1 staining for all tested conditions. The graph illustrates that all permeabilization conditions show good EEA1 antibody staining. Blue shaded bars are to facilitate the evaluation relative to control. Mean ± SEM are displayed. The scale bars of full images are 20µm and inset images are 5µm.

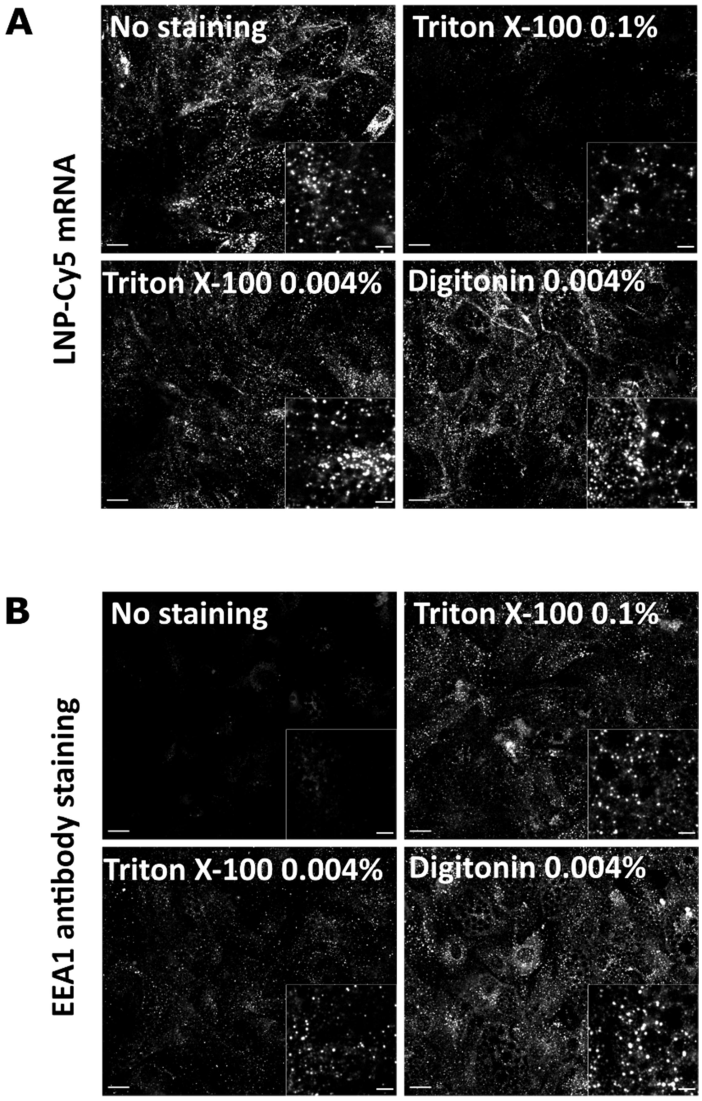


**Supplementary Figure 3: Representative images of mRNA retention and EEA1 staining in human primary adipocytes after Triton X-100 and Digitonin permeabilization.** Experimental details are described in Figure 3C. (**A**) Representative images of LNP Cy5 mRNA signal presented in Figure 3C. The images show Cy5 mRNA signal retained under Triton X-100 and Digitonin permeabilization (0.004%, 1min) condition is comparable to control (no staining). In contrast, the classical method (Triton X-100 0.1% 10min) shows poor retention. (**B**) Representative images of EEA1 antibody staining presented in Figure 3D. Digitonin (0.004%) permeabilization which preserves most mRNA signal also displays good EEA1 staining, whereas Triton X-100 permeabilization (0.004% 1min) yields poor EEA1 staining compared to the classical method (0.1% Triton X-100, 10min). The scale bars of full images are 20μm and inset images are 5μm.

A

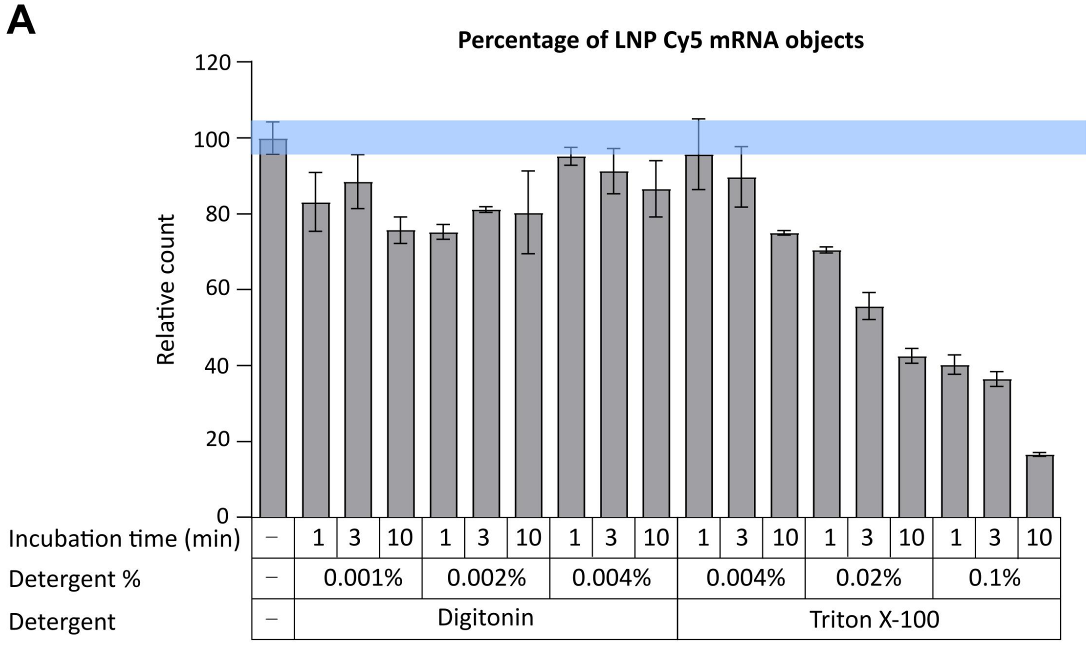


B

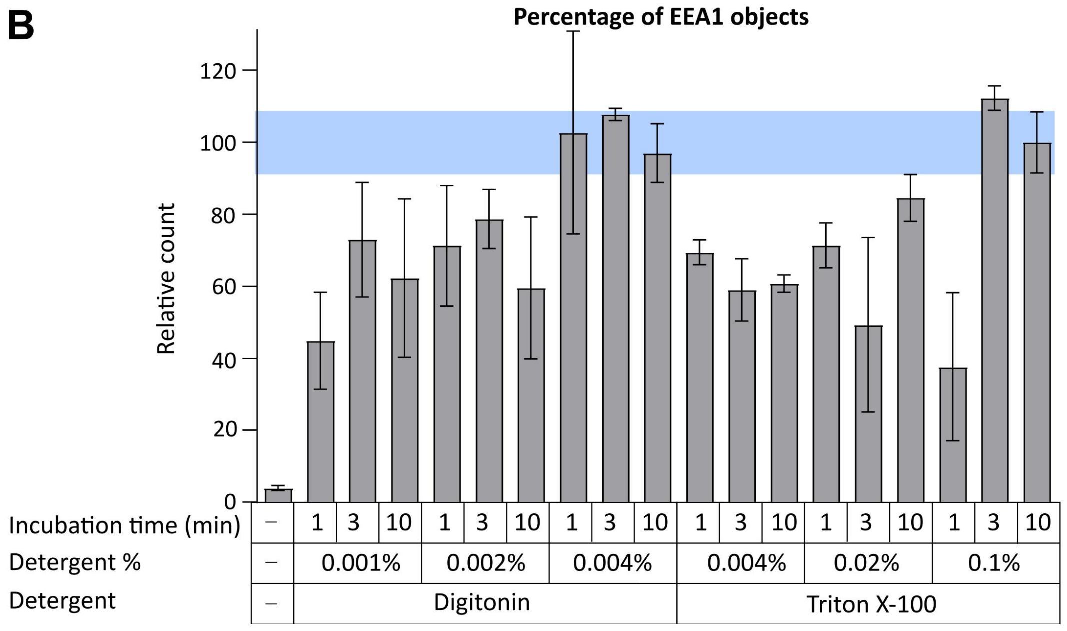


**Supplementary Figure 4: Quantification of mRNA retention and EEA1 staining in human primary adipocytes after Triton X-100 and Digitonin permeabilization.** Experimental details and few selected conditions are described in Figure 3C and D. The first bar in both graphs represents the control of no-staining condition. (**A**) LNP Cy5 mRNA quantification for all tested conditions. The graph shows that Digitonin permeabilization retains most mRNA signal at all concentrations and incubation times. In contrast, Triton X-100 permeabilization shows poor retention with increasing concentrations and increasing incubation times. (**B**) Quantification of EEA1 staining for all tested conditions. The graph illustrates that Digitonin permeabilization at 0.004% yields good EEA1 antibody staining. In contrast, Triton X-100 permeabilization at lower concentrations and shorter incubation times shows poor EEA1 antibody staining. Mean ± SEM are displayed. Blue shade bar is to facilitate the evaluation relative to control.

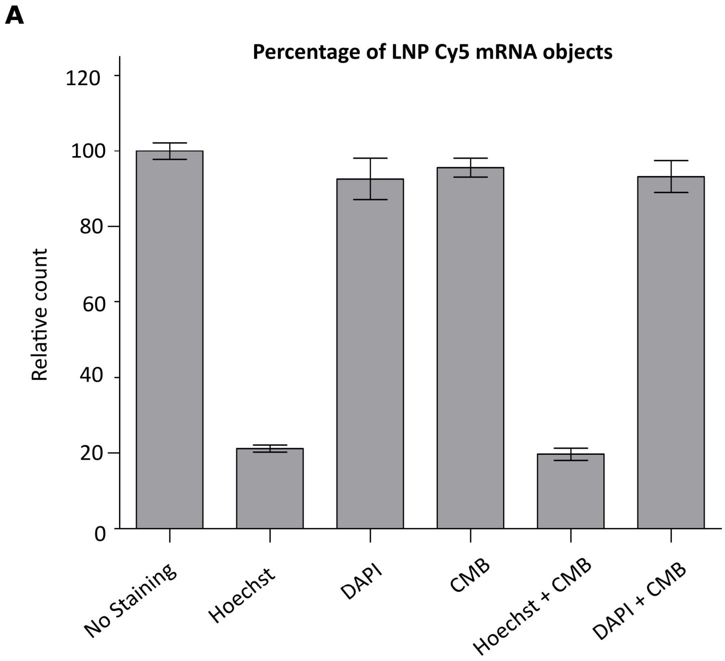


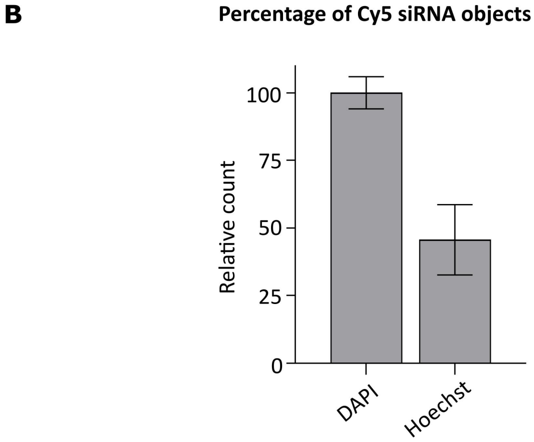


**Supplementary Figure 5: Hoechst staining quenches Cy5 mRNA and siRNA signal.** (**A**) HeLa cells treated with LNP Cy5 mRNA (1h) were fixed with FA (3.7%, 10min) and stained either with DAPI (1µg/mL) or Hoechst (6µg/mL) or CMB (0.25µg/mL) or in combination. The quantification of Cy5 mRNA objects shows that Hoechst staining quenches about 80% of the Cy5 mRNA signal. Experiments were performed in quadruplicates. Mean ± SEM are displayed. B) HeLa cells treated with Interferrin+ Cy5 siRNA (10nM, 30min) were fixed and stained as stated above for Hoechst and CMB. Similar to mRNA, the siRNA signal is quenched about 55%. N = 3 independent experiments. Mean ± SEM are displayed.

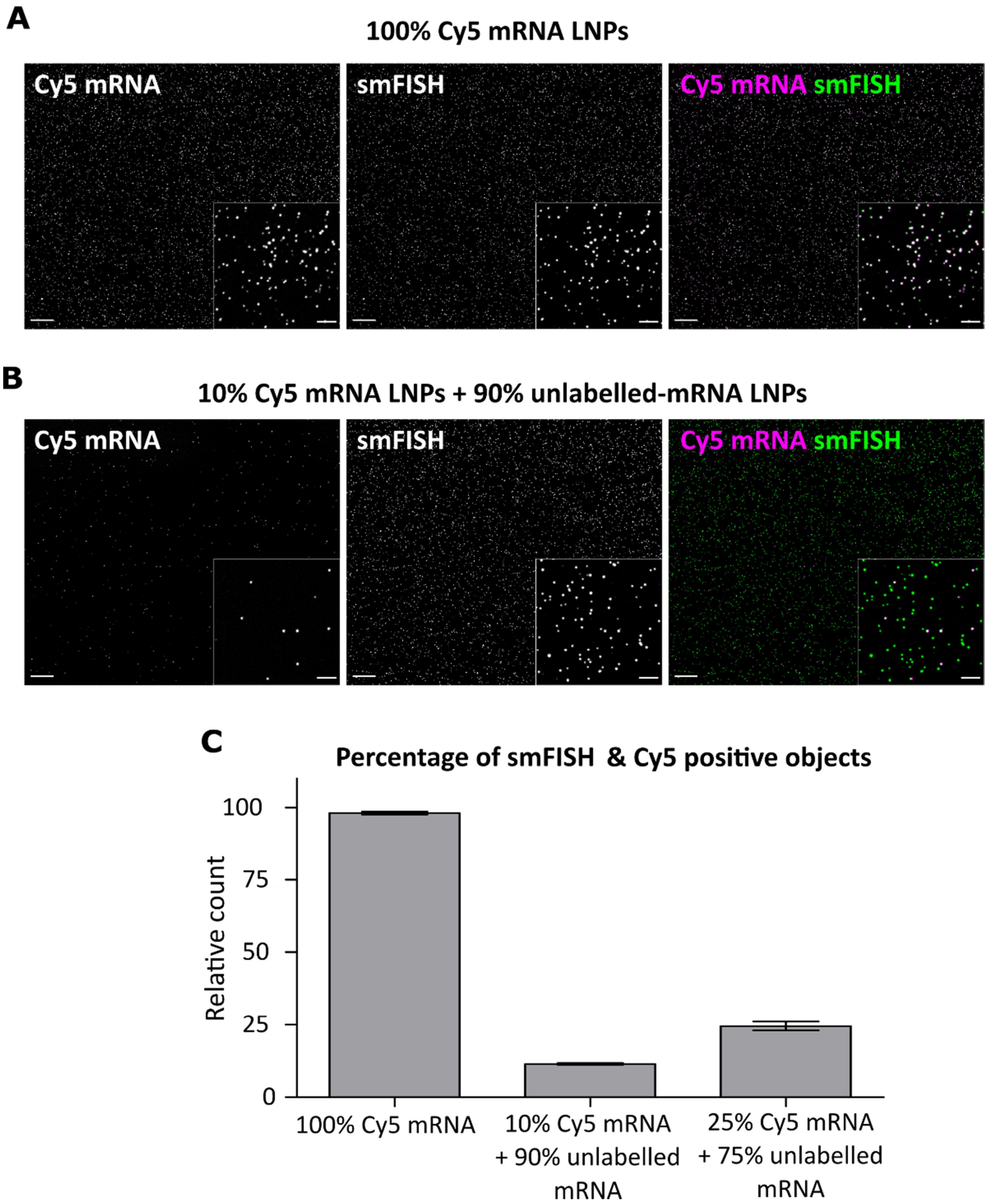


**Supplementary Figure 6: smFISH staining of LNP Cy5 mRNA deposited on glass slides:** (**A & B**) LNPs formulated with Cy5 mRNA or unlabeled mRNAs were mixed at indicated ratios, added into wells without cells (30min), fixed (7.4% FA, 2h) and performed smFISH. The percentage of smFISH and Cy5 double positive objects directly co-relates to percentage of Cy5 mRNA LNP present in the sample. (**C**) Quantification of Cy5 mRNA and smFISH double positive objects. The graph shows the percentage of smFISH and Cy5 double positive objects co-relate with the amount of Cy5 mRNA present in the sample. This indicates that smFISH labelling of Cy5 mRNA is quantitative and efficient. Mean ± SEM are displayed. The scale bars of full images are 20µm and inset images are 5µm.

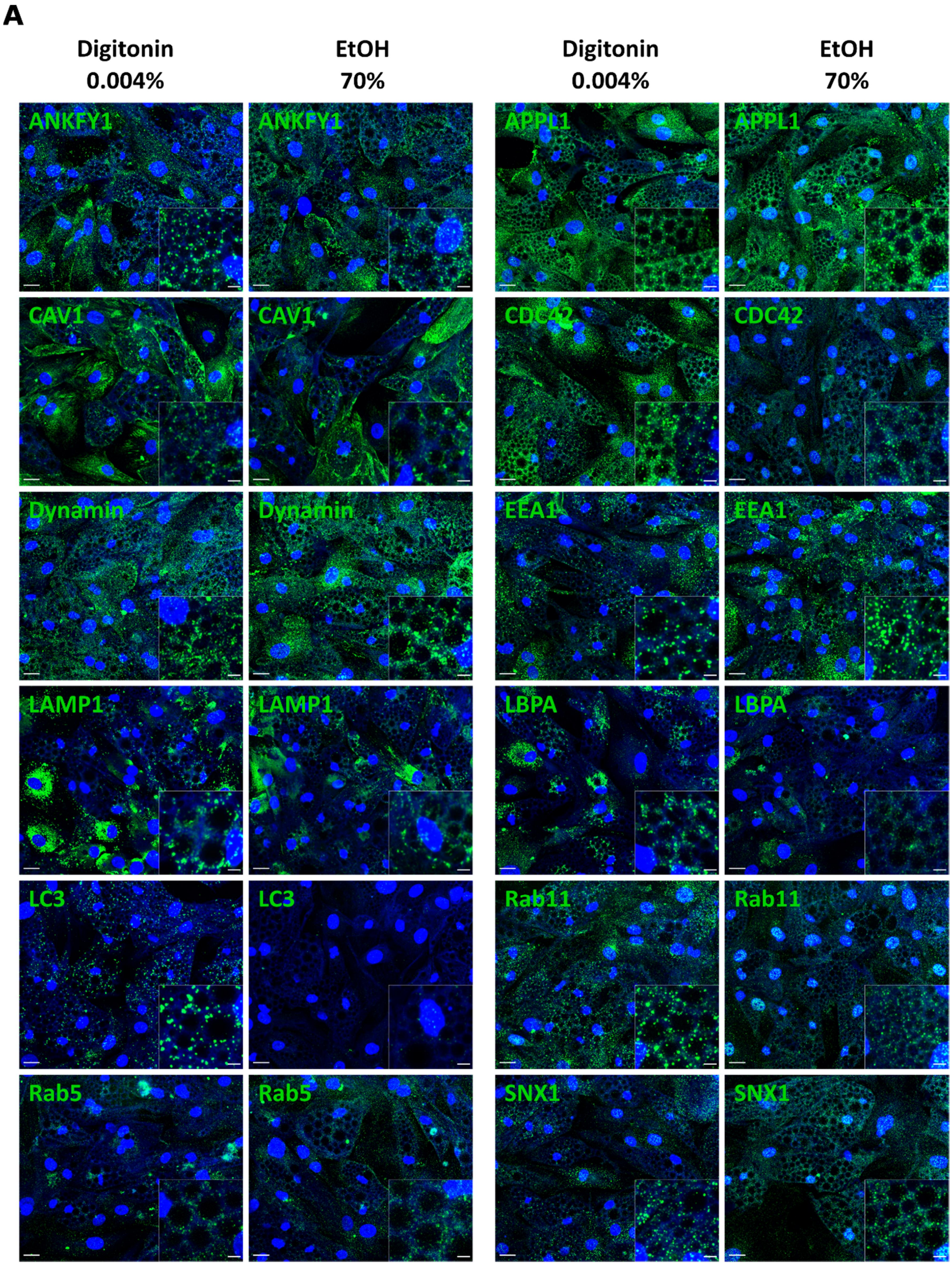


**Supplementary Figure 7: Represetantive images of various antibody stainings under Digitonin and ethanol permeabilization.** (**A**) Cells were fixed with 7.4% FA for 2h and permeabilized with 0.004% Digitonin or 70% ethanol. IFS under ethanol permeabilization shows poor staining for LC3, CDC42, LBPA and Rab11 antibodies. Additionally, there are frequent nuclei stainings using APPL1, Rab11 and SNX-1 antibodies. The scale bars of full images are 20µm and inset images are 5µm.

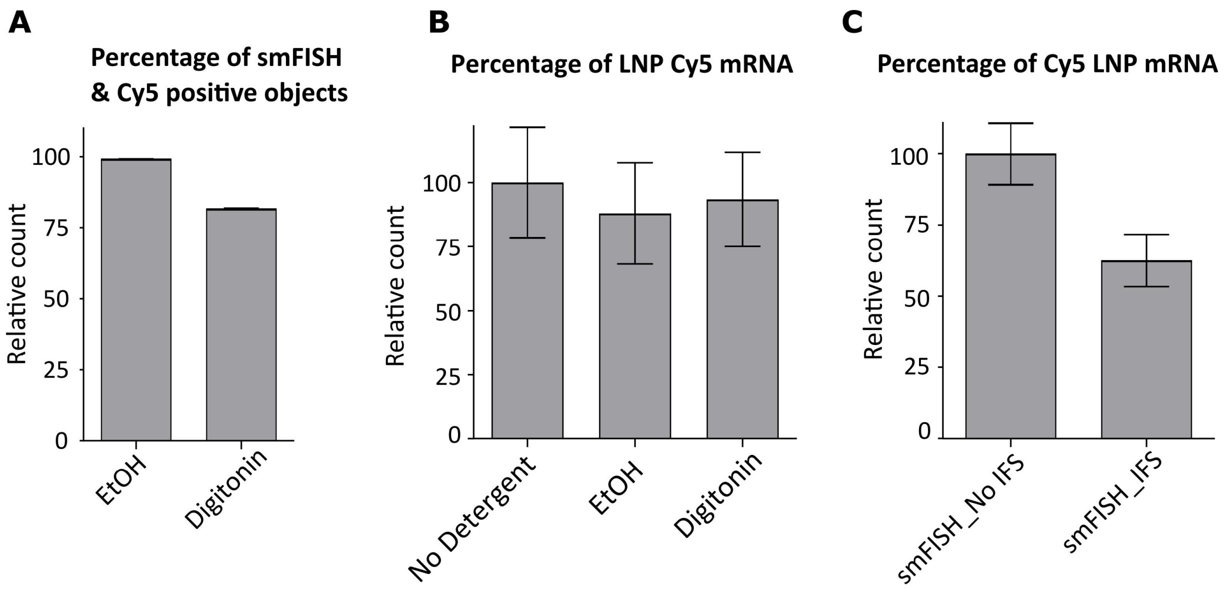


**Supplementary Figure 8: Adaptation of improved fixation and IFS methodolgy to smFISH in HeLa cells.** (**A**) Quantification of Cy5 mRNA and smFISH double positive objects shows that the smFISH labelling of Cy5 mRNA is efficient in cells permeabilized with ethanol, where as a subset of Cy5 mRNA objects are not labelled in Digitonin-permeabilized cells. (**B**) Quantification of Cy5 mRNA-positive objects in cells permeabilized with ethanol or Digitonin. The graph illustrates that Cy5 mRNA positive objects are retained after Digitonin and after ethanol permeabilization. (**C**) The graph shows the percentage of Cy5 mRNA positive objects. About two thirds of the Cy5 mRNA signal was retained after IFS and smFISH staining. Mean ± SEM are displayed.

A

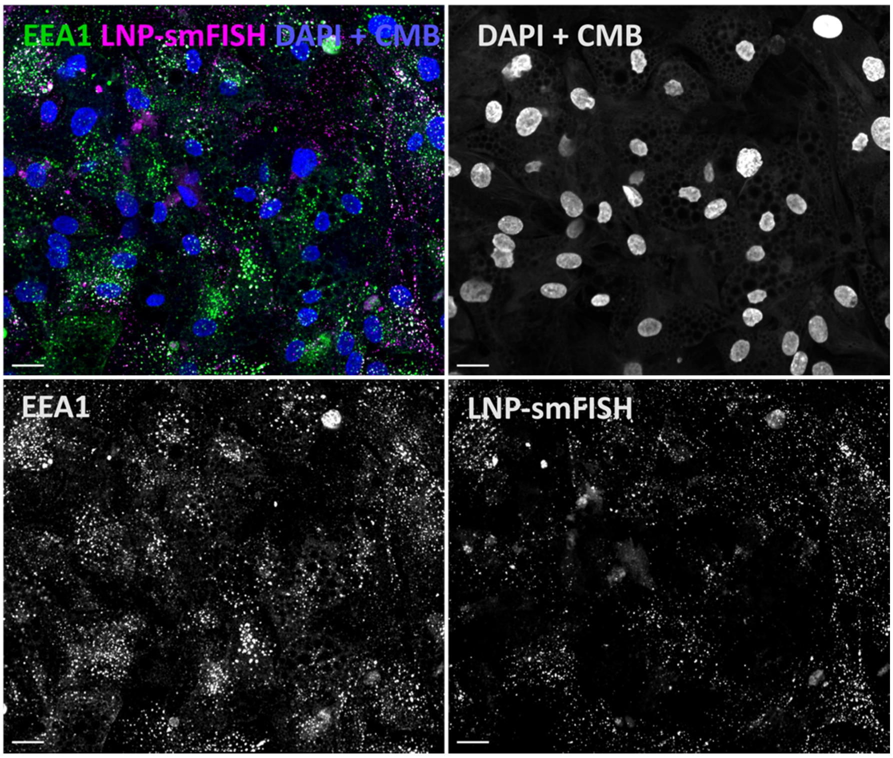


**Supplementary Figure 9: smFISH and IFS in adipocytes using improved methodology.** (**A**) The experimental details are provided in the result section. The images show smFISH stained mRNA (LNP-smFISH) and EEA1 antibody staining in human primary adipocytes using improved methodology. The scale bars are 20µm.

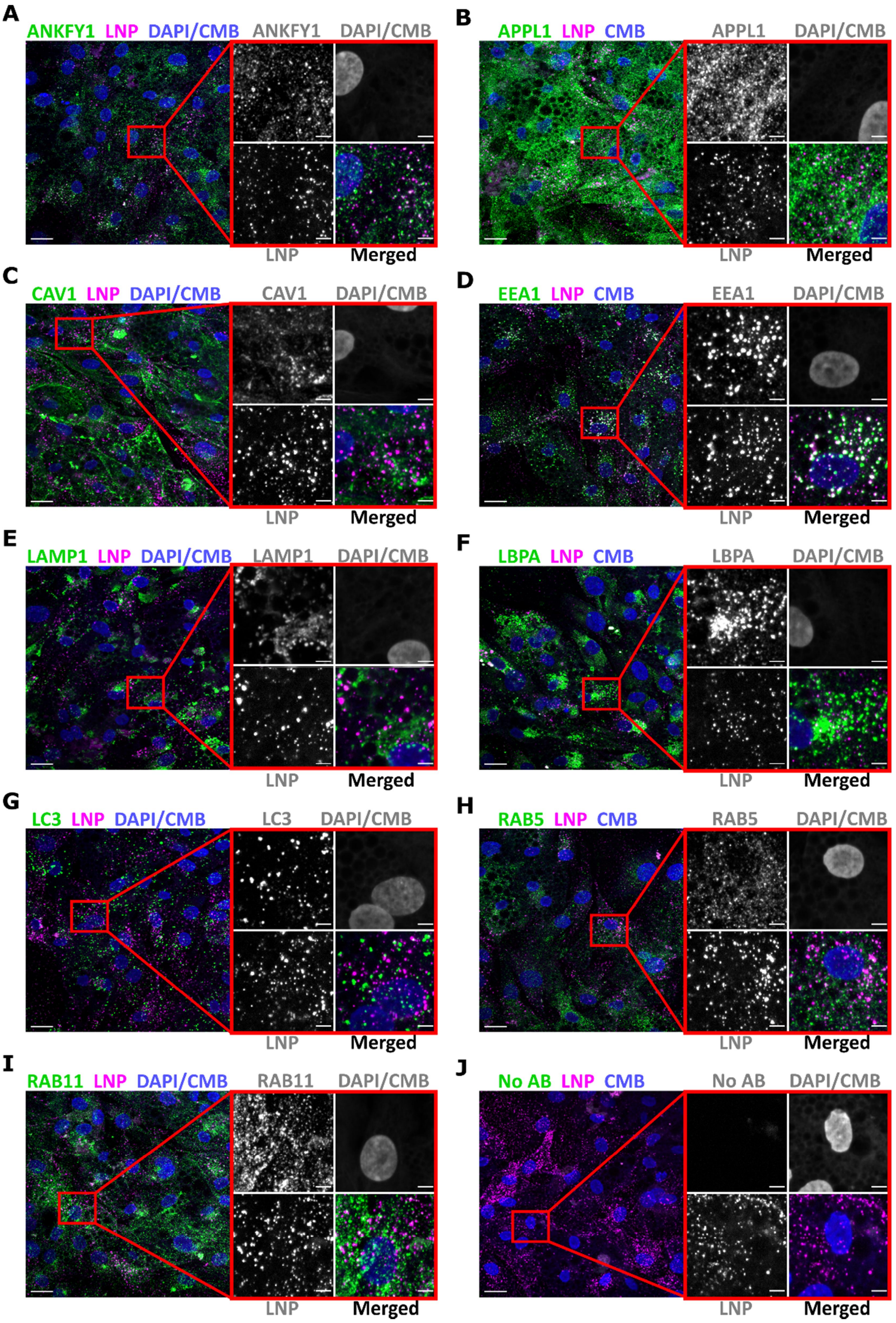


**Supplementary Figure 10: Improved method is compatible with several markers of endosomal compartments.** (**A-J**) Representative images of adipocytes stained for endosomal compartments and exogenously delivered mRNA. Cells incubated with LNP unlabeled mRNA (1h) were fixed (7.4% FA, 2h), performed IFS to mark indicated endosomal compartments and smFISH to detect exogenously delivered mRNA as described in the main text. Zoom-in regions are presented with split and merged color. The scale bars are 20µm in the overview and 5µm in the zoom-in images.

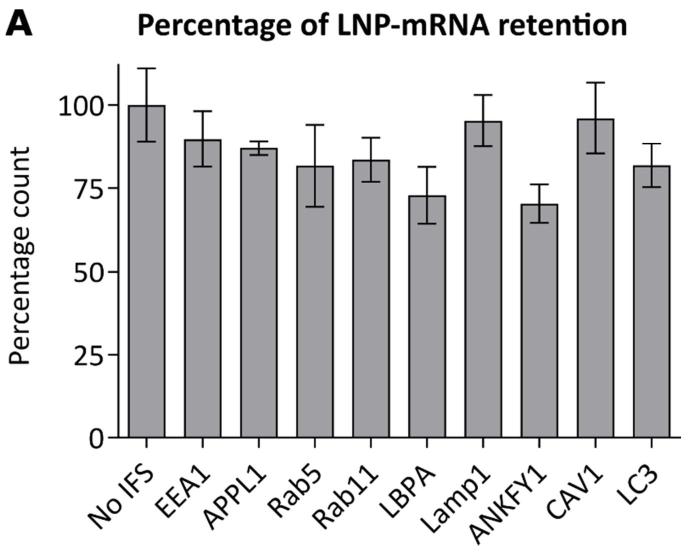


**Supplementary Figure 11:** (**A**) The quantification of mRNA retention after IFS and smFISH shows that mRNA signal retention is consistent. 3 replicates per condition, Mean ± SEM.

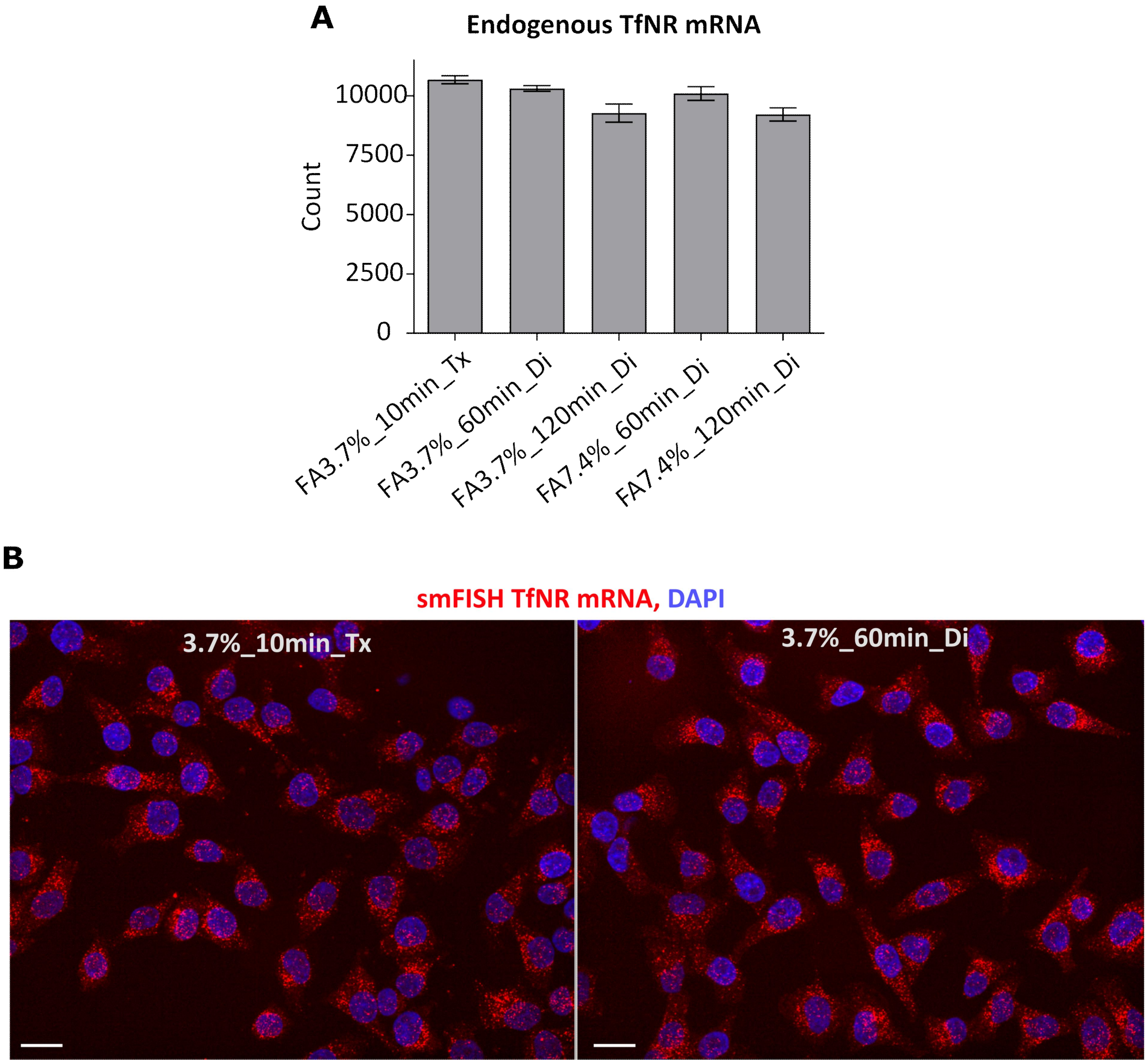


**Supplementary Figure 12: smFISH of endogenous mRNA with improved methedology.** The graph show smFISH stained transferrin receptor endogenous mRNA (TfNR mRNA) in HeLa cells using improved methodology. (**A**) Quantification of TfNR mRNA show no mRNA loss after Triton-X 100 (Tx) 0.1% 10min compared to Digitonin (Di) 0.002% 2min permeabilized conditions under various fixation times and concentrations of FA. (B) Representative images smFISH TfNR mRNA under 10min 3.7% FA fixed and Triton X 100 permeabilized cells vs 60min 3.7% FA fixed and Digitonin permeabilized condition. The scale bars are 20µm.

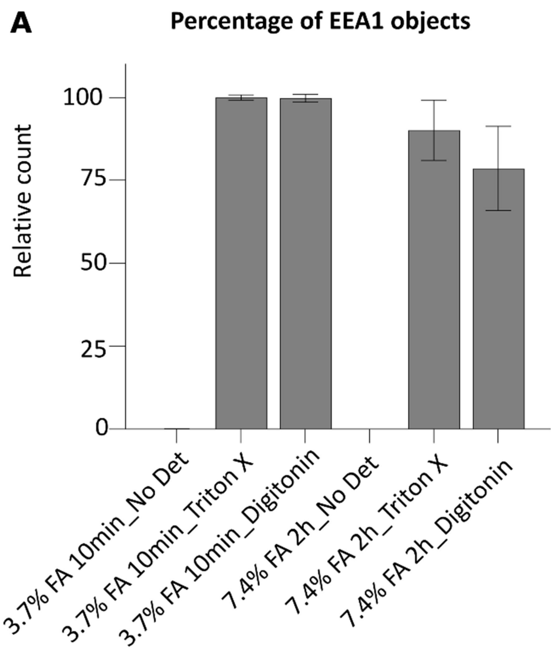


**Supplementary Figure 13: EEA1 antibody staining under high concentration and longer fixation time in HeLa cells:** (**A**) The graph shows that under 7.4% FA 2h fixation, EEA1 signal is reduced. Data collected from the same experiments presented in Figure 6B. N = 3 independent experiments. Mean ± SEM are displayed.

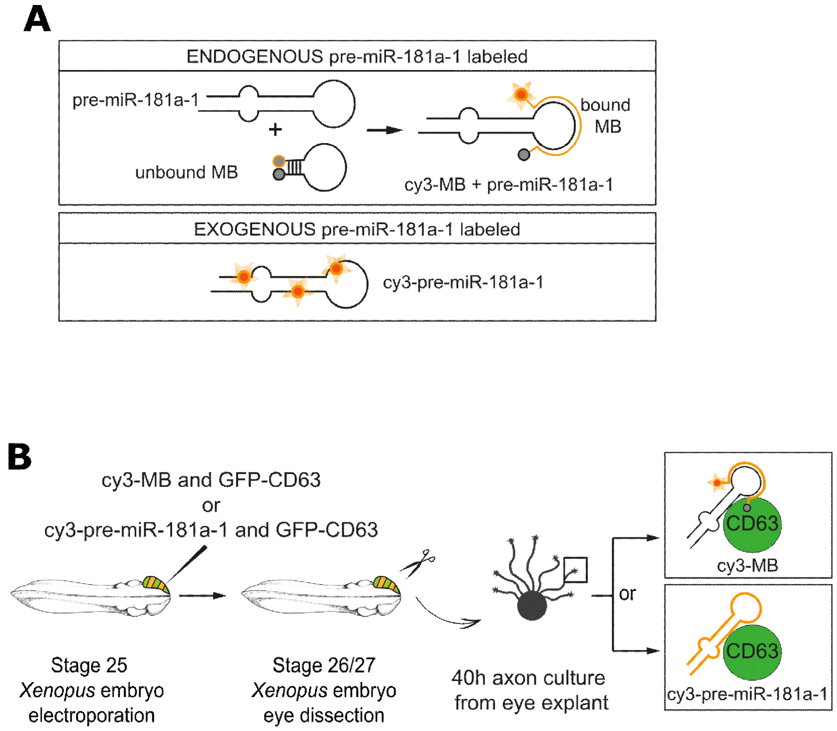


**Supplementary Figure 14:** (**A-B**) **Experimental overview of miRNA labeling and detection from *Xenopus* organoculture.** Details are described in the main text.

| Antibody name | Company | Product number | Antibody dilution used |
|---|---|---|---|
| **ANKFY1** | Sigma Aldrich | SAB1401696-50UG | 1 to 100 |
| **CAV1** | Cell Signaling | #3238 | 1 to 100 |
| **SNX-1** | Thermo Fisher Scientific Inc. | PA1-21544 | 1 to 100 |
| **Lamp1** | BD Bioscience | 555798 | 1 to 200 |
| **LC3** | MBL | M152-3 | 1 to 500 |
| **Rab5** | BD Transduction Laboratories Bioimaging | 610725 | 1 to 100 |
| **APPL1** | Produced at Eurogentec, Belgium (1). Purified at MPI-CBG | α APPL1 2624-3 | 1 to 250 |
| **Dynamin** | BD Bioscience | 610246 | 1 to 100 |
| **EEA1** | Produced at EMBL (2). Purified at MPI-CBG | α EEA1 f.1 Ø7JF | 1 to 1000 |
| **LBPA** | Echelon Biosciences/ MoBiTec | Z-SLBPA | 1 to 100 |
| **Rab11** | Invitrogen (Thermo Fisher) | 71-5300 | 1 to 250 |
| **Donkey polyclonal anti rabbit Alexa Fluor 647** | Invitrogen (Thermo Fisher) | # A31573 | 1 to 1000 |
| **Donkey polyclonal anti mouse Alexa Fluor 647** | Invitrogen (Thermo Fisher) | # A31571 | 1 to 1000 |

**Supplementary Table 1: Details of antibodies and their dilutions used in adipocyte and HeLa cell experiments.**

Miaczynska, M., S. Christoforidis, A. Giner, A. Shevchenko, S. Uttenweiler-Joseph, B. Habermann, M. Wilm, R. G. Parton and M. Zerial (2004). "APPL Proteins Link Rab5 to Nuclear Signal Transduction via an Endosomal Compartment." Cell **116**(3): 445-456.

Simonsen, A., R. Lippe, S. Christoforidis, J.-M. Gaullier, A. Brech, J. Callaghan, B.-H. Toh, C. Murphy, M. Zerial and H. Stenmark (1998). "EEA1 links PI(3)K function to Rab5 regulation of endosome fusion." Nature **394**(6692): 494-498.
